# Decoding Cytokine Networks in Ulcerative Colitis to Identify Pathogenic Mechanisms and Therapeutic Targets

**DOI:** 10.1101/2024.09.12.612623

**Authors:** Marton Olbei, Isabelle Hautefort, John P. Thomas, Luca Csabai, Balazs Bohar, Hajir Ibraheim, Aamir Saifuddin, Dezso Modos, Nick Powell, Tamas Korcsmaros

## Abstract

Ulcerative colitis (UC) is a chronic inflammatory disorder of the gastrointestinal tract characterised by dysregulated cytokine signalling. Despite the advent of advanced therapies targeting cytokine signalling, treatment outcomes for UC patients remain suboptimal. Hence, there is a pressing need to better understand the complexity of cytokine regulation in UC by comprehensively mapping the interconnected cytokine signalling networks that are perturbed in UC patients. To address this, we undertook systems immunology modelling of single-cell transcriptomics data from colonic biopsies of treatment-naive and treatment-exposed UC patients to build complex cytokine signalling networks underpinned by putative cytokine–cytokine interactions. The generated cytokine networks effectively captured known physiologically relevant cytokine–cytokine interactions which we recapitulated in vitro in UC patient-derived colonic epithelial organoids. These networks revealed new aspects of UC pathogenesis, including a cytokine subnetwork that is unique to treatment-naive UC patients, the identification of highly rewired cytokines across UC disease states (IL22, TL1A, IL23A, and OSM), JAK paralogue-specific cytokine-cytokine interactions, and the positioning of TL1A as an important upstream regulator of TNF and IL23A as well as an attractive therapeutic target. Overall, these findings open up several avenues for guiding future cytokine-targeting therapeutic approaches in UC, and the presented methodology can be readily applied to gain similar insights into other immune-mediated inflammatory diseases (IMIDs).

**One Sentence Summary:** A systems immunology map of cytokine interaction networks in ulcerative colitis reveals novel insights into disease pathogenesis, with potential to guide future cytokine-targeting therapeutic strategies.

## INTRODUCTION

Cytokines are key signalling molecules of the immune system that play critical roles in immune response regulation. Cytokines are released by a wide variety of cell types, and can bind to their cognate receptors on the surface of their target cells (1). The cytokine ligand binding to its receptor initialises a signalling cascade, culminating in the regulation of downstream target genes, including other cytokines (2). Thus, cytokines can form complex networks of interactions, in which different cytokines regulate the production and activity of other cytokines, thereby organising immune responses. Many of these individual cytokine interactions have been well established in classical immune responses, such as IFNG activation by IL12 as part of the Th1 adaptive immune response (3, 4), which in turn activates chemokines such as CXCL10 through innate immune cell types such as monocytes (5). Such cytokine–cytokine interactions are important for maintaining the proper balance and specificity of immune responses. However, when dysregulated, chronic immune-mediated inflammatory disorders such as ulcerative colitis (UC) can develop (6). UC is a major subtype of inflammatory bowel disease, characterised by chronic colonic inflammation limited to the mucosal layer of the gut wall. UC causes significant morbidity to patients with symptoms such as bloody diarrhoea, abdominal pain, and fatigue, and its prevalence is rising globally (7). UC pathogenesis is underpinned by an imbalance of pro- and anti-inflammatory cytokine production, leading to dysregulated cytokine signalling, chronic inflammation, and ultimately tissue damage (8–10).

Many of the currently available biologic therapies in UC aim to target key dysregulated cytokines. These include anti-TNF agents such as infliximab and adalimumab, biologics targeting the common p40 subunit of IL12 and IL23 such as ustekinumab, and more recently drugs targeting the p19 subunit of IL23 such as mirikizumab. In addition, small molecule drugs inhibiting JAK signalling (such as filgotinib, tofacitinib, and upadacitinib) have been established in clinical practice in recent years, which underpin multiple cytokines and immune signalling cascades (11). The addition of these advanced therapies in the therapeutic armamentarium has significantly increased the prospects for UC patients, resulting in better outcomes and quality of life (12). However, despite this, a substantial therapeutic ceiling exists, in which 50-70% of the patient population fails to respond or become resistant to these treatments (13–15). This has led experts to call for novel therapeutic strategies targeting multiple cytokines of interest, such as combination biologic therapies. Indeed, clinical trials such as the VEGA trial (evaluating anti-TNF and anti-IL23 combination therapy) are currently underway to test this hypothesis (16). As such, there is an increased need to understand the complexity of cytokine regulation in IBD, to disentangle how treatment impacts cytokine signalling, and how this information can be leveraged to offer alternative perturbation points (17, 18). However, systems-level analyses collecting and modelling the totality of these cytokine–cytokine interactions have been few and far between (19, 20).

In our previous work, we established CytokineLink, a novel systems biology framework that maps cytokine–cytokine interactions based on ligand and receptor expression (19). Although CytokineLink allows for the exploration of these interactions, it lacks integration of the intracellular signalling cascades that follow cytokine-receptor binding. In this study, we substantially advance our network models by incorporating intracellular signalling data derived from single-cell gene expression profiles of UC patients (21), enhancing the accuracy and depth of cytokine–cytokine interaction analysis. Using a novel integrated pipeline, we reconstructed condition-/ treatment-specific cytokine networks of the colonic mucosa in UC patients and non-IBD controls. Comparative analysis of these cytokine networks captured important cytokine regulation differences between treatment-naive UC patients, treatment-exposed UC patients, and non-IBD controls. The analysis identified the cytokines with the most altered interactions in UC, a unique cytokine subnetwork in treatment-naive patients, and novel insights into cytokine signalling hierarchies. It also highlighted the pervasive role of JAK signalling in cytokine interactions and the specificity of JAK paralogues in UC. Overall, this systems immunology approach provides unique insights into UC pathogenesis with significant translational potential, and can be employed to delineate cytokine signalling in other immune-mediated inflammatory diseases (IMIDs).

## RESULTS

### Mapping the cytokine interaction networks driving UC

To better understand how cytokines may regulate each other in UC, we generated cytokine–cytokine interaction networks from single-cell RNA sequencing data derived from colonic biopsies of UC patients and non-IBD controls. To achieve this, we utilised the recently published single-cell RNA-Seq compendium for IBD, scIBD (21), which provided us curated, batch-corrected, and integrated scRNA-Seq data from multiple studies, improving the power of our analysis. We used the data from 85 UC patients and 43 non-IBD controls from scIBD, in total. scRNA-Seq data from UC patients were stratified based on the inflammation status of the colonic biopsies and further grouped into immunotherapy (immunomodulators or advanced therapies) naive or treated patients using the available supplementary materials of the original studies.

We harnessed the computational methods of NicheNet to infer ligand-target gene interactions from the scRNA-Seq data (22). This approach incorporates a prior model regarding intracellular signalling and gene regulatory networks to predict if the binding of ligands to target receptors can modulate the expression of target genes. This was important to capture the potential intracellular mechanisms connecting upstream and downstream cytokines, and mapping the chain of events consisting of cytokine → receptor → signalling proteins → transcription factors → cytokine genes (Figure 1A). As we focussed on cytokine–cytokine interactions, we restricted both upstream ligands and downstream target genes to known cytokine genes as defined in established systems immunology databases. This allowed us to construct networks of cytokine–cytokine interactions from the scRNA-Seq data. For further details on network construction, please refer to the Methods section.

**Figure 1.**
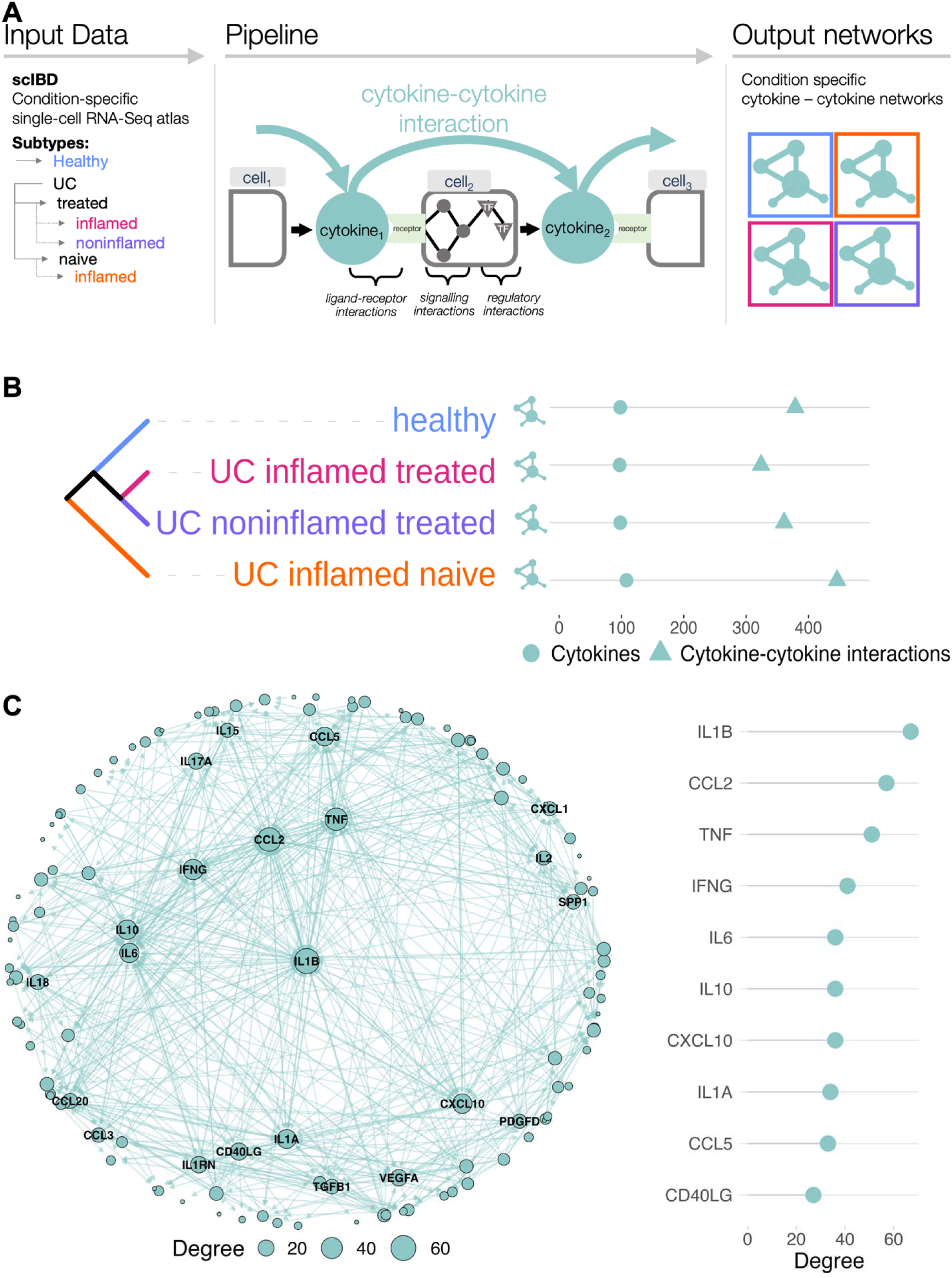
Generating condition-specific cytokine networks in ulcerative colitis. **A**: The computational pipeline processes data from the single-cell RNA-Seq inflammatory bowel disease atlas scIBD. Based on available metadata, we delineated a novel cluster of inflamed UC samples (“inflamed-naive” in the above figure) from patients who did not receive any immunomodulatory or advanced therapies. Data was processed using NicheNet, with the gene set of interest set to a list of cytokines. It connected active cytokine ligands to their putative cytokine targets, resulting in condition-specific cytokine networks. **B**: Hierarchical clustering of networks according to disease and treatment status. The generated networks are comparable in size, with the inflamed treatment naive network having the largest number of interactions. **C**: Analysing all generated interactions reveals the highest degree cytokines in the cytokine networks. The centrality network layout emphasises betweenness centrality, with the most central cytokines positioned closer to the centre.

The datasets within the scIBD atlas differentiate the UC subtypes based on inflammation status, containing samples from both inflamed and noninflamed locations of the gastrointestinal tract. To add further dimensionality to the generated networks, we delineated a novel cluster of biologic treatment-naive inflamed UC samples from the publicly available supplementary materials of the included datasets. Hierarchical clustering of the generated cytokine networks revealed that the networks cluster according to disease status (UC vs non-IBD control), treatment exposure, and inflammation status (Figure 1B). For each condition, the generated networks were comparable in size and shared 43% of their interactions across all states.

The local and global importance of nodes was assessed using degree (i.e., local importance, number of neighbours) and betweenness centrality (i.e., global importance), respectively. Including interactions from all cytokine networks, the highest degree cytokines are IL1B, CCL2, TNF, IFNG and IL6 (Figure 1C). The same five cytokines also achieved the highest betweenness centrality measures (albeit in slightly different order: IL1B, CCL2, IFNG, TNF, IL6), highlighting not only their local, but also their global importance in cytokine communication.

### Cytokine networks capture physiologically relevant cytokine–cytokine interactions

To confirm the physiological relevance of the cytokine networks, we tested our results with a large independent compendium focusing on immune cells and with our organoid experiment focusing on UC patient-derived epithelial cells. First, we compared our data to the recently published Immune Dictionary resource (23). Briefly, the Immune Dictionary encompasses the responses of 17 murine immune cell types to 86 cytokines one-by-one. We tested the target overlap and set similarity of each matching source (i.e. upstream) cytokine from our cytokine–cytokine interaction networks (Figure 2A). 45 cytokines cause differential expression of other cytokines in the Immune Dictionary resource. Out of the 45, 20 cytokines are also source nodes in our networks, targeting other cytokines. Comparing their sets of targeted cytokines with the UC cytokine network model, 12 individual cytokines have a significant overlap in their targets between the Immune Dictionary results and our computational model, 8 do not (60%) (hypergeometric test, FDR <= 0.05). Out of the remaining 25 comparable cytokines with the Immune Dictionary resource, 17 are present as nodes in our cytokine networks as targets of other cytokines (but not as source molecules targeting other cytokines). 8 cytokines were not among our search parameters. The significant overlap between key targets in our constructed cytokine–cytokine interaction networks and those in the Immune Dictionary resource suggest that our networks are likely to be physiologically relevant. However, we acknowledge not all interactions could not be captured due to inherent differences in the sets of cytokines between both studies - for example, the present research involves many chemokines and growth factors that were not included in the Immune Dictionary resource. Furthermore, the cellular compartments of the producing cells also differed, as the Immune Dictionary primarily focussed on immune cell types from lymph nodes, while the UC cytokine networks presented in this study were generated from colonic tissue samples from patients involving the immune, epithelial and stromal compartments. Finally, the Immune Dictionary was generated using murine tissues, and important differences exist between the mouse and human immune systems (24–26).

**Figure 2.**
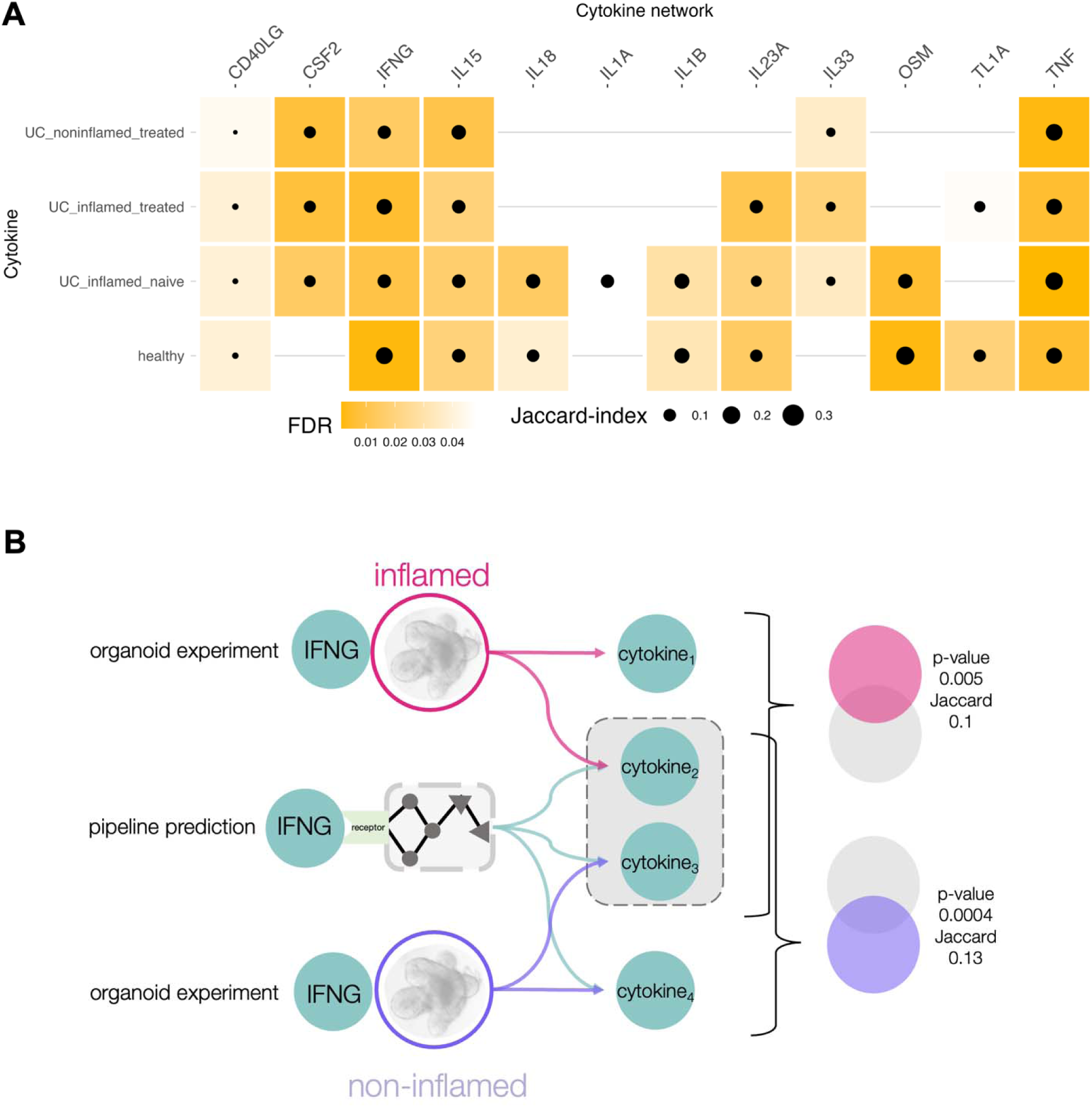
Validation of cytokine–cytokine interactions. **A**: Comparison of target sets between predicted interactions and differential expression results from the Immune Dictionary resource (Cui et al) 12/20 shared source cytokines have significantly overlapping target sets. **B**: Comparison of target sets in epithelial cells between predicted interactions from a reduced computational model only containing cell types also present in colonic organoids, and differentially expressed genes from patient-derived inflamed and non-inflamed colonic organoids (n = 4) treated with IFNG. Set similarity was measured using the Jaccard-index.

To further validate our results, we generated inflamed and non-inflamed colonic epithelial organoids from UC patients (n = 4) to model inflammation and recapitulate the inflamed and non-inflamed interactions in vitro. Organoids were treated with a cocktail of pro-inflammatory mediators (IL1B, TNF, and Flagellin) previously shown to reproduce a similar transcriptomic phenotype to the inflamed epithelium in IBD patients to create inflamed organoids (27). We then treated both inflamed and non-inflamed organoids with IFNG. This proinflammatory cytokine plays a crucial role in intestinal inflammation, it is released by adaptive immune cells, and is known to act on the intestinal epithelial layer (28). The effects of IFNG stimulation on these organoids were captured using RNA-Seq. We compared the overlap of differentially expressed genes from these experiments with a simplified UC cytokine network, which includes interactions exclusively between epithelial cell types relevant to organoids (Figure 2B). We found a significant overlap in the differentially expressed genes responding to IFNG treatment and the target cytokines predicted by the pipeline utilising only epithelial cell types commonly found in organoids (hypergeometric test, inflamed p-value = 0.005, noninflamed p-value = 0.0004). For further details please refer to the Methods section.

### A specific cytokine subnetwork distinguishes treatment naive and treatment exposed inflamed UC tissues

Leveraging the supplementary materials of the original studies from the scRNA-Seq datasets in scIBD, we generated a cytokine–cytokine interaction network from inflamed biopsies of seven treatment-naive UC patients i.e. individuals who had not been exposed to immunomodulators or advanced therapies (for sample IDs and dataset details, please refer to the Methods section). Thus, the network we constructed from these patients likely reflects the nature of cytokine interactions in active UC prior to therapeutic intervention. We determined the Jaccard index for each interaction based on the cell types involved, indicating the similarity of compared sets, in order to determine whether there are differences in the putative cell types mediating these cytokine interactions. We established this in both the treatment-naive and treatment-exposed patient groups, compared to non-IBD controls. Figure 3A highlights the distribution of these scores in the inflamed treatment-naive and inflamed treatment-exposed networks in UC, with the inflamed treatment-naive interactions exhibiting a lower median Jaccard index. This indicates that the cell types contributing to these inflamed treatment-naive interactions are more different than in the inflamed treatment-exposed networks compared to the healthy state (median inflamed-naive = 0.27, median inflamed-treated = 0.38, Wilcoxon-Rank Sum test P < 1.6e-06). There are 17 cell types shared between inflamed-treated and healthy conditions that are not present in the inflamed-naive condition (Supplementary Figure 1).

**Figure 3.**
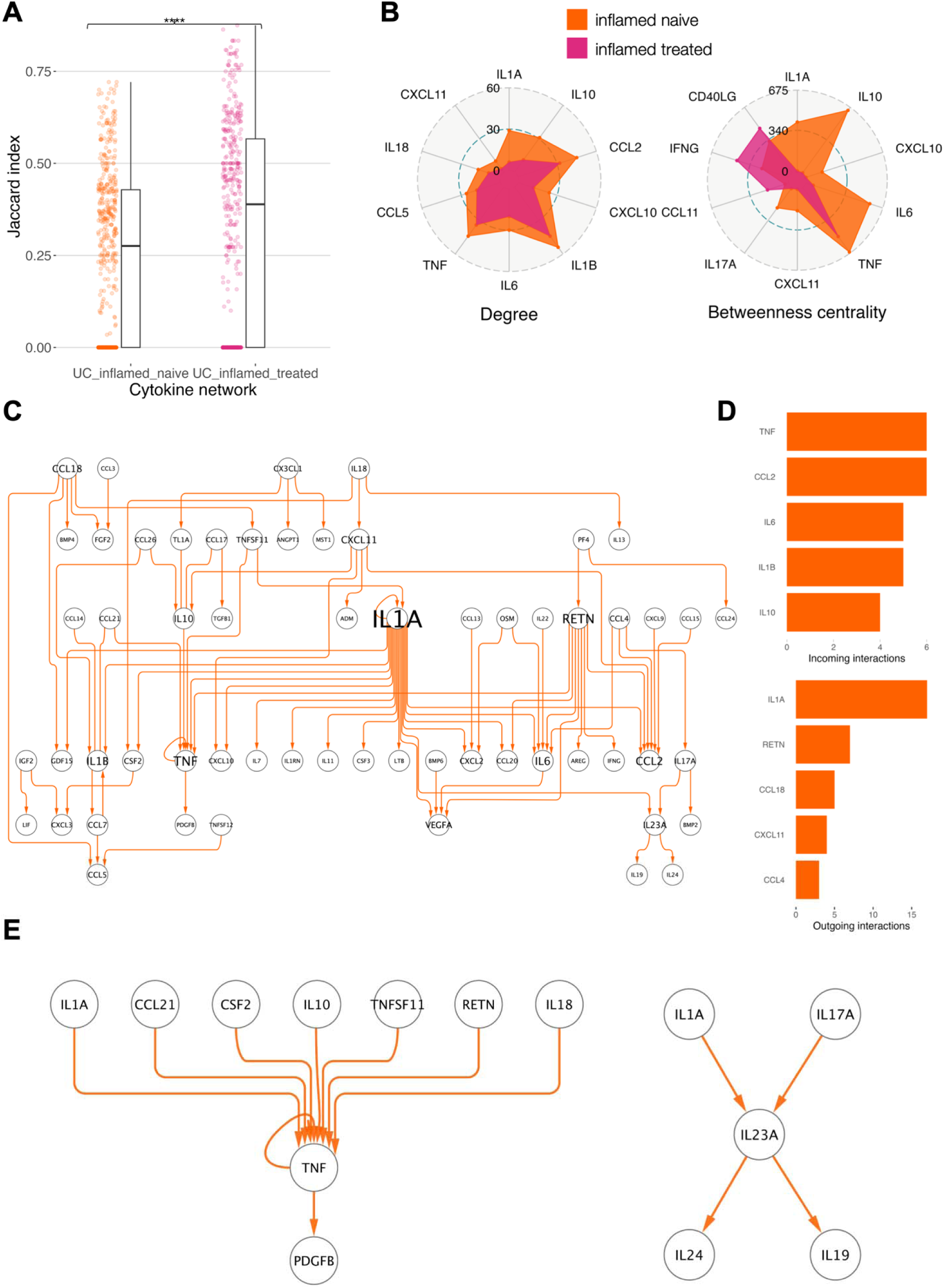
A specific subnetwork of cytokine-cytokine interactions characterise treatment-naive inflamed UC tissues. **A**: Differences in the Jaccard index distribution of cytokine– cytokine interactions. The Jaccard index quantifies the set similarity of cell types contributing to the cytokine–cytokine interaction (as source or target cells) with the healthy condition. The inflamed treatment-naive network has a lower average Jaccard index than the inflamed treatment-exposed network, indicating that the contributing cell types are more different compared to healthy than in the treated networks (median inflamed-naive = 0.27, median inflamed-treated = 0.38, Wilcoxon-Rank Sum test *P* < 1.6e-06). **B**: Degree and betweenness centrality differences between inflamed treatment-naive and treatment-exposed networks. The highest degree cytokines all lose interactors in the treatment-exposed condition, and the highest betweenness centrality cytokines become less central, except CD40LG, IFNG and CCL11. **C**: An inflamed treatment-naive condition-specific subnetwork in ulcerative colitis. **D**: Cytokines with the highest number of incoming interactions in the treatment-naive subnetwork (top) and cytokines with the highest number of outgoing interactions (bottom). **E**: Cytokines with active signals currently targeted by biologic agents (TNF, IL23A) in the the treatment-naive state

We next sought to understand the differences in cytokines mediating these interactions during active disease in the treatment-exposed and -naive groups. We compared the number of neighbours (degree values) of the cytokines present in the networks between both conditions (Figure 3B). We observed that following exposure to immunomodulators or advanced therapies, all cytokines lost interactions with their immediate neighbours, diminishing their local importance. A comparison of their global importance, assessed through betweenness centrality, showed a similar trend, with the exception of three cytokines (CD40LG, IFNG, and CCL11), which became more central after treatment. These findings suggest that exposure to immunomodulators or advanced therapies restructures cytokine networks in UC, reducing both the local and global significance of key cytokines that mediate cytokine–cytokine interactions in active disease.

To better understand the cytokine–cytokine interactions underpinning active UC without any modulation from immunomodulators or advanced therapies, we visualised the interactions that were unique to the treatment-naive group. 83 interactions were unique to the UC inflamed treatment-naive network, representing 19% of the total interactions in this network. 77 of these interactions formed a connected subnetwork. We visualised these treatment naive-specific interactions using a hierarchical layout, allowing us to determine the chain of cytokine interactions involved in the treatment-naive inflamed colonic mucosa. Figure 3C shows the hierarchical structure of these state-specific interactions, and the largest sources and sinks unique to the condition. The highest in-degree nodes of this subnetwork are the important pro-inflammatory cytokines TNF, IL1B and IL6, and the chemokine CCL2 (Figure 3C lower right). The most promiscuous source cytokine is IL1A, targeting 17 other cytokines, highlighting its important role in inflammation (29, 30) (Figure 3C lower right). IL1A is followed by RETN (resistin), an inflammatory marker in IBD with known roles in the inflammatory response (31, 32). Thus, our systems immunology cytokine modelling captured prototypical cytokines associated with UC pathogenesis, whilst providing novel insights into their hierarchical relationships.

We next focussed on cytokines that are currently directly targeted by biologics in UC. The most commonly used biologic agents in UC are anti-TNF agents (33). We found that TNF is one of the highest degree nodes in the treatment-naive inflamed UC cytokine network, interacting with 37 other cytokines i.e. 34% of the cytokine network members. Figure 3D highlights the UC-specific inputs from multiple proinflammatory and regulatory cytokines targeting TNF, with the incoming CCL21, IL10, IL1A, RETN, TNFSF11 interactions only appearing in the UC treatment-naive condition. As for outgoing interactions of TNF, TNF targets itself and PDGFB in the treatment-naive inflamed condition - the latter cytokine previously shown to be an important driver of fibrosis in IBD (34) (Figure 3E). The cytokine IL23A is another important target of biologic treatment in UC (35), either via drugs (e.g., ustekinumab) targeting its p40 subunit shared with IL12A (not active in network), or more recently via drugs targeting its p19 subunit (e.g., mirikizumab). IL23A has four interactions unique to the treatment-naive state: upstream inputs from IL1A and IL17A, and downstream targets to the IL10 family cytokines, IL19 and IL24 (Figure 3E). As with most other cytokines (Figure 3B), the degree and betweenness centrality values for IL23A are reduced in the treatment-exposed inflamed condition.

### Network rewiring analysis pinpoints cytokines with the most varying interactomes across inflammatory states in UC

To characterise the distinct and shared properties of the generated cytokine networks, and to identify both divergent and conserved cytokine interactions across states, we initially analysed the networks using the DyNet network rewiring algorithm (36). This approach allowed us to identify the cytokines present in all networks which have the most variable neighbourhoods (Figure 4A). The top-ranking rewired cytokines (Figure 4B) include IL22, TL1A (encoded by TNFSF15), IL23A and OSM which have been implicated in the pathogenesis of IBD (11, 14, 37)). Figure 4C shows the detailed interactomes of IL22 and OSM.

**Figure 4.**
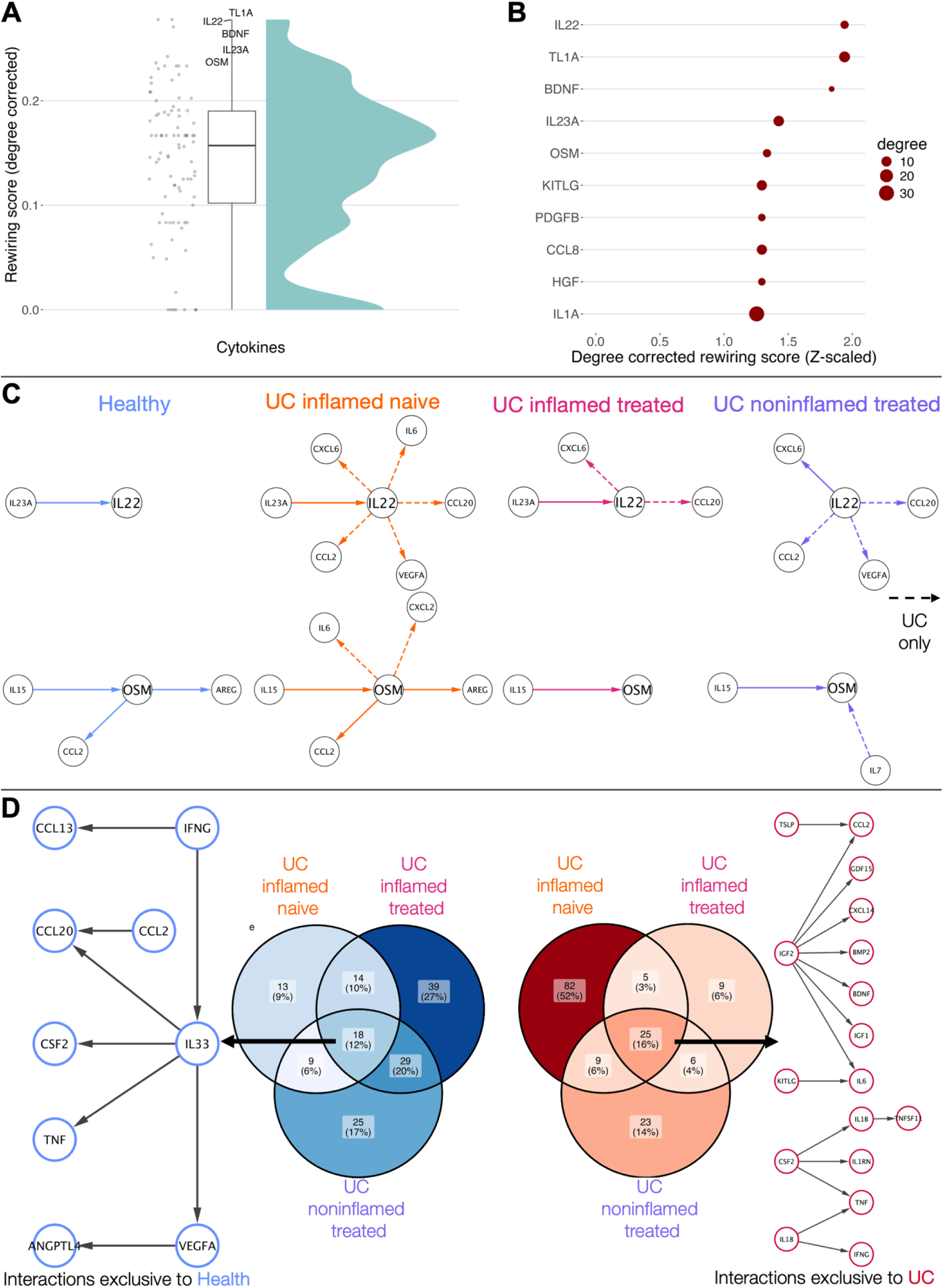
Cytokine network rewiring and shared interactions. **A**: Distribution of degree-corrected rewiring scores. The top-scoring cytokines are highlighted. **B**: The top rewired nodes shared across all networks **C**: In-depth look at the first neighbours of rewired nodes IL22 and OSM. **D**: Missing (left) and shared (right) subnetworks in UC. The largest connected components of the missing (exclusive to healthy) and shared (exclusive to UC) interactions are shown next to the Venn-diagrams.

IL22 is an IL10-family cytokine involved in epithelial regeneration (38). Its levels are often elevated in the intestinal mucosa of IBD patients (11). IL22 has been linked with anti-IL12/23 treatment resistance, and studies have shown its ability to elevate the levels of proinflammatory CXC-family chemokines (37). In the inflamed naive cytokine network, IL22 is targeted by IL23A (39) and can directly regulate CXCL6 and, through CCL2, reach CXCL1-5 (Supplementary Figure 2). It sends further direct UC-specific interactions to IL6, CCL20, and VEGFA.

OSM (oncostatin M) is an IL6-family cytokine whose levels are often elevated in IBD patients (40). OSM is a driver of inflammatory processes, shown to increase the expression of IL6 and proinflammatory chemokines (14), as captured by our model. OSM has also been linked with anti-TNF treatment resistance (14). Besides targeting the aforementioned proinflammatory cyto-/chemokines, OSM also targets AREG in the treatment naive cytokine network, a cytokine shown to promote intestinal fibrosis in experimental colitis and human Crohn’s disease patients, a different subtype of IBD (41).

Following the rewiring analysis, to explore the shared properties of the UC networks, as opposed to their differences, we collated the interactions missing from all UC cytokine networks compared to the healthy state (i.e. interactions exclusive to health), and collected interactions present in all UC cytokine networks (i.e. interactions exclusive to UC) (Figure 4D). There are 18 health-specific, and 25 UC-specific interactions, regardless of treatment status or whether the sample is coming from an inflamed or non-inflamed site. Figure 3D shows the largest connected components of these shared interactions. The central node in the graph of health-specific interactions is IL33, an important IL1-like alarmin responsible for initiating a response to cellular damage (42). So far, there have been conflicting reports regarding the role of IL33 in UC, as it has shown to exert both detrimental and protective effects in mouse models of colitis (43).

In contrast, the shared UC-specific interactions form two larger subgraphs. In the first, the upstream nodes are IL18 and CSF2, cytokines shown to stimulate and, in some cases, decrease the levels of proinflammatory cytokines such as IFNG, TNF and IL1B (11, 44–47). The second connected component is driven by IGF2, primarily targeting other growth factors, many of which were previously implicated in IBD pathogenesis (48–51).

### JAK specificity of cytokine–cytokine interactions in UC

We next evaluated the intracellular pathways mediating cytokine–cytokine interactions in UC, specifically focusing on the JAK signalling cascade which serves as the primary signal transduction pathway for multiple cytokines (52). JAK signalling has emerged as a crucial therapeutic target in IBD following recent phase III clinical trials demonstrating the efficacy of non-selective and selective JAK-inhibitors for induction and maintenance therapy in UC (53). To deconvolute the specificity of JAK paralogues in cytokine signalling in UC, we captured the likely intracellular paths connecting source cytokines to their putative target cytokine genes, and collated the JAK proteins underpinning each interaction. In the UC inflamed treatment-naive network 55% of interactions signalled using one or more JAK proteins (Figure 5A). To assess the reliability of the captured cytokine-JAK relationships, we compared the number of correct and incorrect JAK paralogue assignments for 9 active cytokine ligands in the networks whose JAK preferences were recently reviewed (11), and the networks contained downstream cytokine interactions from them. In 9 out of 9 cases, the correct JAK paralogues were assigned to the signalling cytokine. However, the pipeline did not assign JAK proteins to intracellular paths for a few interactions between certain cytokines (mean 0.72% ± 0.27%) (Figure 5B). To quantify the impact of JAK paralogue-specific inhibition in UC, we calculated the DyNet rewiring score of the UC inflamed treatment-naive cytokine network before and after removing cytokine-cytokine interactions which signal through each affected JAK paralogue (Figure 5C). Based on the UC inflamed treatment-naive cytokine network, the neighbourhoods of CD40LG, IL10, and IFNG, were the most perturbed by the removal of JAK1 mediated interactions, indicating that JAK1 selective inhibitors such as upadacitinib may inhibit cytokine-cytokine interactions connecting to these cytokines. The removal of JAK1-affected edges also excluded certain nodes from the cytokine network, such as OSM and IL22, which have been recently associated with treatment resistance to anti-TNF (14) and anti-IL12/23 (37) therapies respectively (Figure 5D). Thus, JAK1 signalling is likely to encompass these resistance proteins, which may explain why JAK1 inhibition with upadacitinib or non-selective JAK inhibition with tofacitinib demonstrated generally consistent efficacy in both biologic naive and biologic exposed UC patients in phase III clinical trials (54, 55).

**Figure 5.**
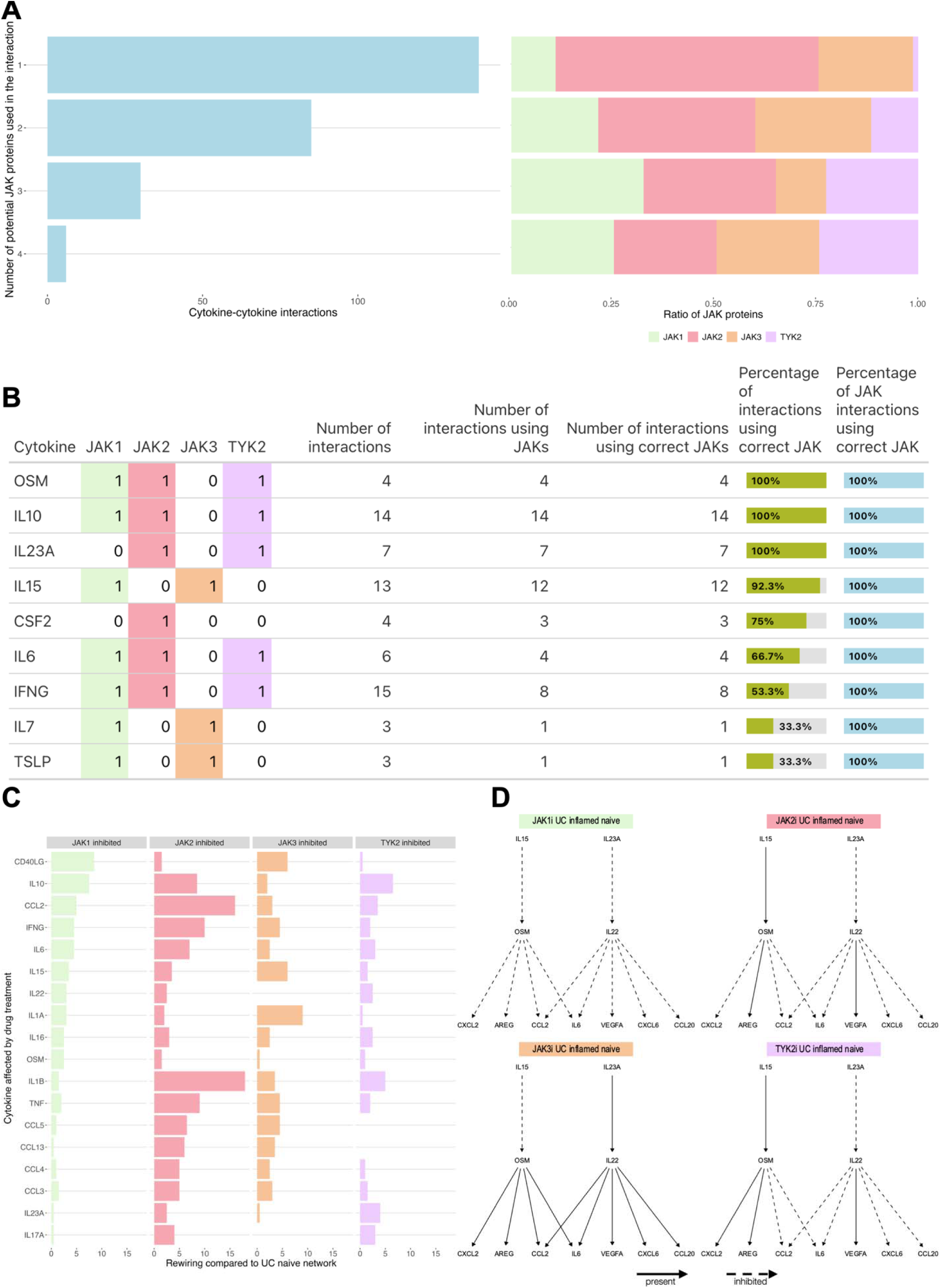
JAK paralogue specificity of cytokine networks. **A**: Number of JAK proteins used by the generated cytokine–cytokine interactions (left) and their ratio of used JAK proteins. **B**: Select cytokines with known JAK preferences, and the percentage of correctly identified JAKs in the predicted intracellular signalling pathways. **C**: DyNet rewiring compared to the inflamed-naive network following the removal of cytokine interactions mediated by JAK paralogues. Only top values are shown. **D**: Removal of JAK1 interactions excludes IL22 and OSM from the UC inflamed treatment naive cytokine network.

### Drug targeting analysis reveals TL1A as a critical cytokine target in UC

TNF, IL23A and IL12A are the three cytokines currently targeted by biologics in UC. To better understand how and why particular cytokines might be better targets than others, we assigned annotation data to the analysed cytokines from the OpenTargets database (56). OpenTargets compiles a global score for each protein assessing its validity as a drug target, based on a collection of evidence from multiple data sources and modalities, such as genetic associations, somatic mutations and RNA expression. In general, there is a moderate positive correlation between the global score in OpenTargets and the degree of cytokines in the UC naive network, indicating that the local importance of cytokines captured by the networks correlates with their candidacy as drugs (R=0.44, P < 1.9e-06, Supplementary Figure 3). Figure 5A highlights the OpenTargets score in the context of cytokine networks analysed in this study. Based on the global score measure of OpenTargets, TL1A (encoded by TNFSF15) is the highest-scoring cytokine target in UC without an approved inhibitor drug. TL1A is a known regulatory cytokine and susceptibility variant in IBD (57–59). It has previously been shown to regulate important proinflammatory cytokines such as TNF, IFNG, IL17A, CSF2, IL6 and IL13 (60), the majority of which TL1A targets directly in the inflamed naive UC cytokine networks. Furthermore, TL1A is also one of the top rewired nodes between the cytokine networks, indicating that its interactome is significantly different across states (Figure 3B). Figure 5C plots the log2 ratio of edge weights and Jaccard indexes of the cytokine–cytokine interactions compared to the healthy condition. To quantify the weight of cytokine–cytokine interactions, we used the number of cell types participating in each cytokine–cytokine interaction (see Methods for further details). The upper right quadrant shows the interactions that not only have low Jaccard scores (dissimilar cell types participating in the interaction) but are also expanded in the UC treatment-naive condition (log2 ratio > 2), indicating that a greater number and variety of cell types signal through TL1A compared to health. In the inflamed treatment-naive network, the TL1A-TGFB1 interaction has one of the lowest Jaccard scores and highest log2 edge weights across all interactions when compared to health, indicating a marked shift in the composition and number of contributing cell types that involve inflammatory fibroblasts and reticular fibroblasts in the inflamed UC condition. As TL1A has also been implicated in fibrosis in IBD (61), this interaction reveals one of the potential mechanisms through which this becomes possible. Figure 6D shows the specific differences in the participating cell types of the TL1A - TNF interaction between health and inflamed treatment-naive UC. Compared to the healthy state, in the inflamed treatment-naive condition there is a substantial expansion in the putative immune cell types that are likely to respond to TL1A to produce TNF including various CD4 and CD8 T cell subtypes, epithelial cell subtypes including Goblet cells, BEST4+ and Tuft cells, and endothelial cells. TL1A sends the most expanded input to TNF compared to all other incoming TNF interactions, indicating the inclusion of additional cell types that produce TNF in response to the TL1A signal. We also noted that TL1A and three of its directly regulated cytokines (CSF2, IL10, IL17A) all target TNF with expanded and disease-specific interactions compared to health (Figures 5E and 5F). TL1A may impact the other important UC drug target, IL23A, through a UC-specific interaction with IL17A (Figure 5E, Figure 5F). Recently, an anti-TL1A drug (PF-06480605) has entered a phase II drug trial (TUSCANY) in ulcerative colitis with positive results (62). The trial demonstrated a significant downregulation of TNF, IL23A and other cytokines (IL1B, IFNG, CCL20) compared to baseline and a reduction in the activity of Th17 and fibrosis pathways. Figure 5F highlights the upstream position of the inhibited cytokine TL1A (blue) in the inflamed treatment-naive network in relation to all cytokines found to be significantly downregulated compared to baseline (yellow) as shown in (62), capturing the potential mechanism through which this regulatory effect occurs, involving a number of UC-specific interactions.

**Figure 6.**
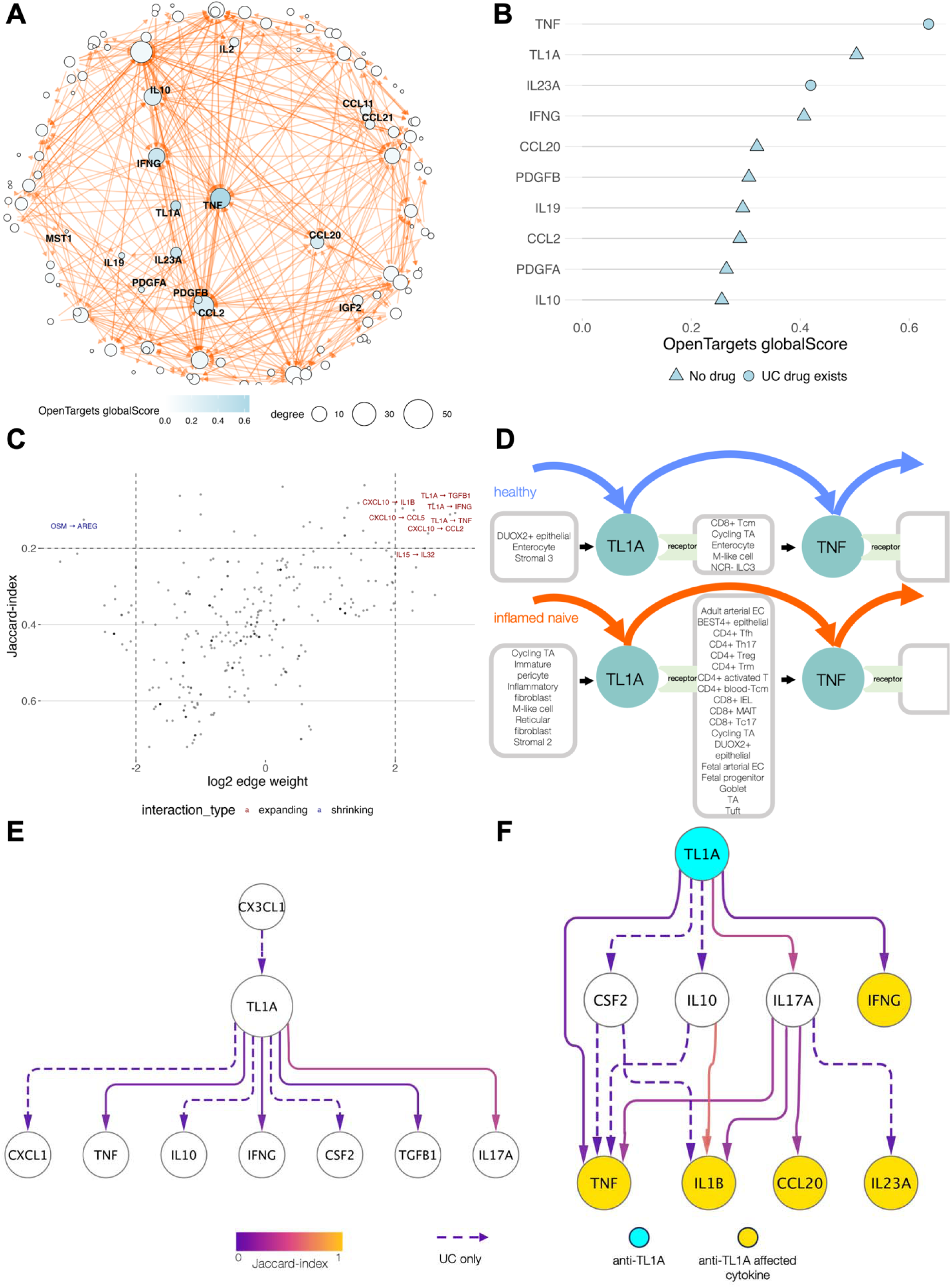
TL1A as a drug target in the treatment-naive cytokine network. **A**: UC inflamed treatment-naive cytokine network with the OpenTargets global score overlaid on the cytokines. **B**: Highest OpenTargets global score of cytokines, and available UC drugs. **C**: Scatterplot of log2 edge weight compared to health and Jaccard index of cell type similarity compared to health. Upper right quadrant indicates interactions that are simultaneously more expanded, and utilise different cell types compared to health. **D**: Cell types contributing to the expanded TL1A-–TNF interaction. **E**: First neighbours of TL1A in the inflamed treatment-naive network. UC-specific interactions shown with dashed lines. **F**: Interactions from TL1A (blue) to all significantly affected cytokines in anti-TL1A treatment as shown in (62).

## DISCUSSION

The complex network of interactions formed by cytokines is an important mechanism regulating immune responses in inflammatory diseases (63). However, the redundant and pleiotropic effects cytokines exert on their target cells make it difficult to disentangle their respective roles (64, 65). In this study, we constructed a novel systems biology model to resolve this by comprehensively collating all active cytokine–cytokine interactions in the system, capturing both the redundant and cell type-specific effects cytokines may have on each other. We demonstrated the ability of this systems immunology model to capture physiologically relevant cytokine-cytokine interactions in both epithelial and immune cell compartments through cytokine exposure experiments in patient-derived colonic epithelial organoids and a large independent compendium of experimentally validated cytokine-cytokine interactions in immune cells (23).

We next sought to gain deeper insights into the cytokine networks underpinning the prototypical chronic immune mediated inflammatory disorder (IMID) of UC across different states based on inflammatory status and treatment exposure. A better understanding of cytokine networks in IMIDs such as UC and how they alter according to inflammatory status or treatment will be critical for finding alternative, and more effective therapeutic strategies (9, 66). In UC, the long-term remission rate of therapeutic agents, including anti-TNF and anti-p40 IL12/23 drugs, remains suboptimal at around 20-30% compared to placebo, imposing a “therapeutic ceiling” (13). For patient cohorts that do not respond well or become resistant to such advanced therapies, finding the cytokine signalling events that differ from responders and/or offer alternative interjection points could help inform strategies for overcoming this therapeutic ceiling in the future.

Our systems biology modelling revealed novel insights into the cytokine networks underpinning UC. We found a large cytokine subnetwork comprising cytokine-cytokine interactions that were specific to treatment-naive inflamed tissues. In this subnetwork, the cytokines with the greatest number of incoming interactions included TNF, IL1B, IL6, and CCL2, whilst IL1A and resistin had the largest number of outgoing interactions. Rewiring analysis of cytokine networks pinpointed cytokines with the most altered interactions across disease states, revealing how treatment and inflammation status can alter cytokine signalling. The most rewired cytokines included IL22, TL1A, IL23A and OSM. We found that TL1A is likely to function as an upstream regulator of key cytokines, including TNF and IL23A, which are already pharmaceutically targeted in UC. Furthermore, many of the cytokines regulated by TL1A involve an increasing number of cell types including CD4 and CD8 T cells, epithelial cells, and endothelial cells in the inflamed UC state compared to the healthy state. We also observed a putative pro-fibrotic signal mediated by TL1A through the activation of inflammatory and reticular fibroblasts resulting in TGFB1 release. Interestingly, TL1A emerged as one of the top druggable cytokine targets in our analysis based on data from OpenTargets. This aligns with results from a recent phase II drug trial in UC targeting TL1A (TUSCANY), which demonstrated reduced expression of TNF and IL23A and other cytokines (IL1B, IFNG, CCL20) captured downstream of TL1A by our model, as well as the alleviation of fibrosis and inflammation in patients following TL1A inhibition (62). Our analysis highlighted multiple other examples where the effect of a cytokine on its downstream targets was also identified in independent studies, such as the effect of IL22 on its target CXC-chemokines (37), the induction of IL22 by IL23A (39), and the targeting of IL6 by OSM (14). Finally, by mapping the JAK paralogue specificity of cytokine-cytokine interactions, including a module involving treatment-resistance genes (OSM, IL22) within the JAK1 signalling network, we revealed a possible mechanistic basis for the consistent efficacy of JAK inhibitors observed in both biologic naive patients and patients who previously failed biologics (54, 55).

The presented systems immunology modelling of cytokine-cytokine interactions allowed us to gain unique insights into UC pathogenesis. Other systems-level models capturing cytokine signalling are rare and tend to answer different questions from those we proposed in our current study. Databases such as ImmuneXpresso and Immunoglobe (67, 68) collate a large amount of cell–cytokine or cytokine–cell interactions compiled from the literature, but do not consider direct cytokine–cytokine interactions, which were the primary focus of our analyses. Another study proposes a method to analyse the effects of the blockage of single cytokines on the secretion rate of target cytokines in IMIDs (20). The approach outlined in the study is a complementary modelling technique that could be combined with our systems-level method to identify the effects on putative intervention targets in the future.

Although our analysis captured physiologically relevant cytokine-cytokine interactions and yielded relevant findings, it is important to note that the presented study has several limitations that must be considered when contextualising the results. The primary focus of our research was to study how cytokines affect each other and observe the rewiring of cytokine signalling in disease. Naturally, cytokines regulate non-cytokine target genes as well, which are not captured in our model, reducing the scope in which the interactions can be interpreted. On the other hand, we used a broad definition of cytokines that includes interleukins, chemokines, growth factors and others to capture a more complete yet potentially less focussed picture of immune signalling. In a complex condition such as UC, where inflammation and wound healing may be constitutively active, we decided on this broad definition to capture a more holistic overview of the underlying pathomechanisms driven by cytokines (69). Importantly, we modelled the individual cytokine–cytokine interactions as direct 1:1 relationships without considering their synergies, an important feature of cytokine regulation (70). However, such synergistic signalling interactions are computationally challenging to model given the lack of comprehensive experimental work that has evaluated multiple cytokines individually and at the same time to delineate such relationships. Another challenge was the limited resolution of clinical meta-data that was publicly available, hindering our ability to identify cytokine signalling changes modulated by specific immunomodulators or advanced therapies. In addition, the interactions can falter on multiple levels in all single-cell RNA-Seq studies focussing on intercellular communication (71). The sequenced RNA in the scIBD datasets might not all translate to protein; if it does, it might not get secreted; and if it does get secreted, it might not diffuse to the target cells, or it might not cause a response (72). Furthermore, cytokine signalling often occurs at an autocrine and paracrine level (73), requiring spatial information which is not captured with single-cell RNA-Seq data. In our model, we have found that over two-thirds of the predicted interactions involve at least one shared cell type producing the primary and secondary cytokines (69% ± 2.2% SD, Supplementary Figure 4), indicating that the majority of cytokine signalling occurs on at least a partially autocrine level. However, with the increasing availability of spatial transcriptomics platforms, there is exciting potential for this limitation to be overcome (74–77).

In the future, more focussed studies are required to further disentangle the hierarchical and signed (stimulatory or inhibitory) nature of cytokine–cytokine interactions. The use of patient-derived organoid models provide a disease-specific genetic background (78), can be co-cultured with certain immune cell populations, and serve as a scalable in vitro model system that can be challenged with multiple compounds (79, 80). Appropriate time series experiments could disentangle whether the cytokine–cytokine interactions are truly directly caused by the addition of an “upstream” cytokine or by second- or third-order effects. While the method discussed in this work uses single-cell RNA-Seq data as its input, alternative methods exist that could be repurposed for cytokine–cytokine interaction inference using less expensive bulk transcriptomics data (81), paving the way to a much bigger repository of publicly available datasets. While more targeted validation is needed to confirm the directionality and stimulatory or inhibitory nature of the predicted cytokine–cytokine interactions, the ability to organise cytokine communication in relevant hierarchies could be harnessed to indicate future upstream targets, if direct intervention would not be available or desirable.

In conclusion, using a novel systems biology approach we mapped the network of cytokine interactions underpinning the chronic IMID of UC revealing distinct interactions according to inflammatory state and treatment exposure. We identified key rewired cytokines across disease states, a subnetwork of cytokine-cytokine interactions in inflamed colonic mucosa unique to treatment-naive UC patients, and pinpointed the putative JAK paralogue proteins participating in each cytokine–cytokine interaction. Our analysis highlighted TL1A as an important upstream regulator of TNF, IL23A, and other cytokines, and a potential driver of pro-fibrotic signalling in UC. Thus, our systems immunology modelling reveals new insights that have the potential to inform the development of novel therapeutic strategies in UC. While this study specifically focused on cytokine signalling in UC, our approach can be used to interrogate cytokine signalling in other inflammatory or infectious diseases to better understand how cytokine interactions contribute to disease pathomechanisms.

## MATERIALS AND METHODS

### Data processing

Integrated, pre-processed single-cell RNA-Seq gene expression data was downloaded from scibd.cn (21) in .rds format. Data were read and processed using the Seurat (82) (version: 4.3) and tidyseurat (83) packages (version: 0.6). Samples were separated by disease condition (i.e., UC, healthy) and status (inflamed, non-inflamed) using the annotation provided by scIBD. Samples annotated as adult tissue biopsies were used for the generation of cytokine networks. In addition to the original disease conditions provided by the scIBD metadata, we created an additional “UC inflamed naive” condition in the ulcerative colitis dataset. Using the supplementary information from the studies included in scIBD, where patient treatment data was assigned to individual samples (rather than on a cohort level). Seven samples (studies: (2) Kinchen et al. (84), (3) Parikh et al. (85), (2) Corridoni et al. (86), sample identifiers: ‘GSM3214201_A3’, ‘GSM3214204_B3’, ‘GSM3214207_C3’, ‘GSM4483695_S24’,’GSM4483696_S33’,’GSM3140595_UC1’,’GSM3140596_UC2’) in ulcerative colitis that did not receive any immunomodulatory or biologics treatment, and thus were considered treatment naive. Subsequently, these patients were removed from the “UC inflamed” category, which we then termed “UC inflamed treated”. We attempted to create similar subclasses for the ‘UC noninflamed’ condition, but in that case only three patients could be assigned to the “UC noninflamed naive” condition, which due to its small size was not comparable with the rest of the data. However, these three patients were excluded from the ‘UC noninflamed treated’ condition to achieve a more consistent output.

### Construction of cytokine–cytokine networks

We used the R package NicheNet (prior model v2) to establish the cytokine–cytokine networks in IBD, for each disease condition (healthy, UC: inflamed naive, inflamed treated, noninflamed treated) (22). The Seurat identity classes of the scIBD object were set “minor_cluster”, containing the established cell types in the dataset. The gene set of interest was determined as all cytokines that were considered such in the systems immunology databases ImmunoGlobe and ImmuneXpresso (67, 68) (Supplementary Table 1). NicheNet’s ligand activity analysis was carried out for each cell type, querying cytokine ligands, indicating their potential in regulating their target gene sets of interest (downstream cytokines). Ligands were considered active if they achieved a Pearson correlation value higher than 0.1. The top-predicted target genes of the active cytokine ligands were established using NicheNet’s target prediction evaluation protocol, a multi-ligand random forest model with k-fold cross-validation (k = 5). Target downstream cytokine ligands from the gene set of interest were kept if they were predicted in every cross-validation round.

### Edge weights

To quantify the weight of cytokine–cytokine interactions we used the number of cell type interactions participating in each cytokine–cytokine interaction. Individual cytokine–cytokine interactions are constructed by connecting cytokine producing cell types with NicheNet, that can plausibly influence each other’s activity. In general terms:

cell1 → cytokine1 → cell2 → cytokine2

Specific examples:

A. , Inflammatory monocyte → IL1B → CD8+ → IL4: an IL1B → IL4 cytokine cytokine interaction
B. , Inflammatory monocyte → IL1B → CD16+ NK → IL4: a different IL1B → IL4 cytokine cytokine interaction, where the site of IL4 production differs.

For the above example the edge weight for IL1B → IL4 would be 2, as two combinations of cell types provide the cytokine–cytokine interaction in this case.

To better highlight differences between the edge weights compared to healthy, we calculated their log2 ratios compared to healthy, e.g., log2FC_IL1B_IL4_inflamed_naive = log2(IL1B-IL4_inflamed_naive / IL1B-IL4_healthy).

### Network rewiring

Cytoscape (version 3.9.1) (87) app DyNet (version 1.0) (36) was used to calculate the rewiring values between the network states, using the multiple comparison mode.

### Network characteristics

The global and local importance of cytokine nodes in the networks was calculated using the tidygraph package (version 1.2.3), using the centrality_degree and centrality_betweenness functions. For the hierarchical clustering of networks the dissimilarity matrix was calculated from the adjacency matrices of the networks using the dist function in R, using the “euclidean” method. Clustering was performed using the hclust function, with the method set to “complete”. The resulting object was written to a newick file using the ape package (88), and visualised using the iTOL web service (89).

### Cytokine paths

To find the shortest paths between queried source and target nodes we used the PathLinker (90) Cytoscape application, with default settings.

### Anti-TL1A affected cytokines

We used the results from Figure 2B of (62) to collect the list of cytokines significantly downregulated following anti-TL1A treatment compared to baseline (IL1B, IL23A, CCL20, IFNG, TNF).

### JAK specificity of cytokine–cytokine interactions

We used the get_ligand_signaling_path function from NicheNet to query the intracellular nodes in each cell type used by each cytokine–cytokine interaction, with the “top_n_regulators” argument set to 10, and the NicheNet weighted network filtered to contain only expressed genes. For each cytokine–cytokine interaction the JAK proteins involved in the signalling paths were recorded. To assess the precision of the affected interactions we compared the number of correct and incorrect JAK usage in 9 cytokines where their JAK preferences are known and downstream targets are found in the network (11).

### Open Targets global score

The OpenTargets global score was downloaded by querying “ulcerative colitis” (EFO_0000729) on the Open Targets Platform (https://platform.opentargets.org/) (accessed 01/02/2024). The results were exported as a .tsv file, and the resulting hits filtered to only contain cytokine genes present in our study. For the calculation of the global score and source data weights please refer to: https://platform-docs.opentargets.org/associations. Correlation with degree was calculated using the stat_cor function of the ggpubr package (version 0.6.0).

### Three-dimensional (3D) organoid culture

Four human organoid lines were used for our study from the Imperial BRC Organoid biobank. These lines were generated from rectal or sigmoid colon tissue from patients with quiescent to mild ulcerative colitis (2 women and 2 men between 30 and 72 years of age). Derived organoid lines were cultured as 3D structures embedded in Matrigel ExtraCellular Matrix, (Corning, Growth Factor Reduced Cat # 356231) diluted with Advanced DMEM/F12 medium using a 4:1 ratio (Thermofisher Scientific, Cat # 12634) in Corning™ 3527 24-well plates (VWR International Gmbh, Cat # 734-1605). Organoids were expanded fed every other day with complete Intesticult Organoid Growth Medium (OGM, Human, StemCell Technologies, Cat # 06010), supplemented with 10µM Rho Kinase inhibitor Y-27632 (Tocris, Cat # 1254) and 100µg/ml Primocin (Invivogen, Cat # ant-pm-1) at 37°C, 5%CO2 in a humidified incubator.

Organoid were then differentiated in complete Intesticult Organoid Differentiation Medium (ODM, Human, StemCell Technologies, Cat # 100-0214) supplemented with 10μM Gamma y-secretase inhibitor (DAPT) (Biotechne Ltd, Cat #2634/10) and 100µg/ml Primocin. Differentiation was also carried out at 37°C, 5%CO2 in a humidified incubator. Organoids were cultured for 4 days prior to any treatment.

### Organoid treatments

Inflammatory status of differentiated UC patient organoids was regained through a 24h treatment with a cocktail containing 100ng/ml TNFα (Invivogen, Cat # rcyc-htnfa), 20ng/ml IL-1β (Peproteck, UK, Cat # 200-1B) and 100ng/ml Flagellin (Invivogen, Cat # tlrl-fliC-10) in complete ODM containing 10μM DAPT and 100µg/ml Primocin.

When required, 100ng/ml IFNγ (Thermo Fisher Scientific (Hemel), Cat # PHC4031) was added to the medium concomitantly or not to the inflammatory cocktail for the 24h incubation prior to the experiment end point.

At the end of the experiment, the potential cytotoxicity of condition media was tested on an aliquot of medium collected from each technical replicate using the CytoTox 96® Non-Radioactive Cytotoxicity Assay kit (Promega, Cat # G1780).

Each condition was tested in 8 technical replicates for each organoid line (biological replicates).

### RNA extraction

At the end of the experiment, each organoid-containing Matrigel dome was rinsed with pre-warmed DPBS (Merck, Cat # D8537). For each condition, organoids were harvested in cell recovery solution (Corning®, Cat # 354253), technical replicates were pooled two by two to ensure enough material is obtained for RNA subtraction. Organoids were then spun down at 300x g, +4°C, for 5 minutes and immediately lysed in the β-Mercaptoethanol-containing lysis solution from the RNeasy minikit (Qiagen Cat # 74104). RNA was extracted following the Qiagen kit instruction manual and eluted in 10mM Tris-HCl, pH8. All samples with an RIN >=7 were used for sequencing.

### Library preparation and sequencing

The libraries for this project were constructed by the Technical Genomics Team at the Earlham Institute, Norwich, UK using the NEBNext Ultra II RNA Library prep for Illumina kit (NEB#E7760L) NEBNext Poly(A) mRNA Magnetic Isolation Module (NEB#E7490L) and NEBNext Multiplex Oligos for Illumina® (96 Unique Dual Index Primer Pairs) (E6440S/L) at a concentration of 10µM. Library preparation was performed on the Perkin Elmer (formerly Caliper LS) Sciclone G3 (PerkinElmer PN: CLS145321). 1µg of RNA was purified to extract mRNA with a Poly(A) mRNA Magnetic Isolation Module. The technology is based on the coupling of Oligo d(T)25 to paramagnetic beads which facilitates the binding of poly(A)+ RNA. Isolated mRNA was then fragmented for 12 minutes at 94°C, and first strand cDNA was synthesised. This process reverse transcribes the RNA fragments primed with random hexamers into first strand cDNA using reverse transcriptase and random primers. The second strand synthesis process removes the RNA template and synthesises a replacement strand to generate ds cDNA. Directionality is retained by adding dUTP during the second strand synthesis step and subsequent cleavage of the uridine containing strand using USER Enzyme (a combination of UDG and Endo VIII). NEBNext Adaptors were ligated to end-repaired, dA-tailed DNA. The NEBNext Adaptors with novel hairpin loop structure are designed to ligate with high efficiency and minimise adaptor-dimer formation. The loop contains a U, which is removed by treatment with USER Enzyme to open the loop and make it available as a substrate for PCR. The ligated products were subjected to a bead-based purification using Beckman Coulter AMPure XP beads (A63882) to remove most of the un-ligated adaptors. Adaptor Ligated DNA was then enriched by receiving 10 cycles of PCR (30 secs at 98°C, 10 cycles of: 10 secs at 98°C _75 secs at 65°C_5 mins at 65°C, final hold at 4°C). Barcodes are incorporated during PCR using NEBNext Multiplex Oligos for Illumina® (96 Unique Dual Index Primer Pairs) thereby allowing multiplexing. The size of the libraries was estimated using the Perkin Elmer GX Touch DNA High Sensitivity assay (DNA High Sensitivity Reagent Kit CLS760672) and the concentrations were quantified by fluorescence, with a high sensitivity plate reader Quant-iT™ dsDNA Assay Kit, (ThermoFisher Q-33120). The resulting libraries were then equimolarly pooled and q-PCR was performed on the pool prior to sequencing. The pool was sequenced on one lane of the Illumina NovaSeq 6000 S4 flow cell with 150 bp paired-end reads.

### Bulk RNA-Seq processing

Bulk RNA-Seq data was preprocessed by our in-house TranscriptOmiX pipeline (https://github.com/korcsmarosgroup/Transcriptomix). The pipeline is made up of the following steps: Pre-alignment Quality Control (FastQC (version v0.12.0) (91), Generating genome index (STAR (version 2.7.10a) genomeGenerate with sjdbOverhang 99, using GRCh38.108 genome annotations), Alignment (STAR alignment) (92), Quantification (featureCounts (version 1.22.2) (93), Post Alignment Quality control (MultiQC (version 1.14) (94). Differential expression analysis was carried out on normalised counts, by DESeq2 (version 1.34.0) (95) (padj_cutoff = 0.05, alpha = 0.05) with lfcShrink, comparing all pairwise conditions.

### Validation of cytokine–cytokine interactions (cytokine–cytokine interactions)

To validate the physiological relevance of cytokine–cytokine interactions, we compared them to two experimental datasets.

First, we downloaded the differential expression results of the Immune Dictionary study from Cui et al. (23) (Supplementary Table 3 of Cui et al.), to compare cytokine responses in immune cell populations to upstream cytokines administered to mice models. Mouse gene identifiers were mapped to human symbols using NicheNet’s convert_mouse_to_human_symbols function. The significance of target set overlap and set similarity were determined between the Immune Dictionary results, and all generated cytokine networks, where comparable source cytokines (i.e. cytokines affecting other cytokines) were found.

To compare the computational results to the organoid experiments, the generated cytokine networks from the UC inflamed treated and UC noninflamed treated conditions were filtered to contain only interactions between epithelial cell types also present in colonic organoid models (Goblet, Enterocyte, Cycling TA, TA, Enteroendocrine, Adult colonocyte, Pediatric colonocyte, BEST4+ epithelial). We compared the downstream cytokine targets of IFNG in these reduced models with differentially expressed cytokine genes from bulk RNA-Seq data from patient derived ulcerative colitis colonic organoids treated with IFNG (for experimental methods see: Organoid culture, IFNG treatment above). Differential expression was established with a |log2FC| >= 2 threshold, and a corrected p-value cutoff of 0.05. Multiple testing was corrected with the Benjamini-Hochberg method.

To calculate the significance of the overlap between cytokine target genes captured by the computational model and the experiments used for validation, for both the Immune Dictionary results and the organoid experiments, we used the hypergeometric test from R (version 4.1.2) with the phyper function, with the argument lower.tail set to false. Multiple testing was corrected using the “fdr” method of the p.adjust function in R. The similarity of the input sets was measured with the Jaccard-index, calculated with a custom R function.

## Acknowledgments

We thank the past and present members of the Korcsmaros and Powell groups, the participants of the Interdisciplinary Signaling Workshop 2023, and Dr. Domenico Cozzetto for their advice and suggestions regarding our work. We thank the NIHR Imperial BRC Organoid Facility for their support of the organoid work, and the Earlham Institute Technical Genomics group for their work involving RNA sequencing.

## Funding

T.K. and I.H. were supported by the NIHR Imperial Biomedical Research Centre Organoid Facility, and the UKRI BBSRC Gut Microbes and Health Institute Strategic Program BB/R012490/1 and its constituent projects BBS/E/F/000PR10353 and BBS/E/F/ 000PR10355, as well as a UKRI BBSRC Core Strategic Program Grant for Genomes to Food Security (BB/CSP1720/1) and its constituent work packages, BBS/E/T/000PR9819 and BBS/E/T/ 000PR9817. B.B, T.K. and I.H. were also supported by the UKRI BBSRC Institute Strategic Programme Food Microbiome and Health BB/X011054/1 and its constituent project BBS/E/F/000PR13631. D.M. acknowledges financial support from Imperial College London through an Imperial College Research Fellowship grant award. A.S. received funding from the International Organisation of Inflammatory Bowel Disease and St Mark’s Hospital Foundation. J.P.T is supported by the Chain-Florey Clinical PhD Fellowship jointly funded by the National Institute for Health Research (NIHR) Imperial Biomedical Research Centre (BRC) and the UKRI Medical Research Council (MRC) Laboratory of Medical Sciences (LMS). N.P. was supported by the Wellcome Trust (WT101159 and 225875). N.P., A.S., and H.I. were supported by the National Institute for Health Research (NIHR) Imperial Biomedical Research Centre (BRC). The views expressed are those of the authors and not necessarily those of the NIHR or the UK Department of Health and Social Care.

## Author contributions

Conceptualisation: MO, TK

Software: MO, LC, BB

Validation: IH, AS, HI

Analysis: MO, JPT, DM

Writing: MO, JPT, IH, DM, TK

Supervision: TK, NP

Funding acquisition: TK, NP

## Competing interests

A.S has received travel expense funding from Janssen and Alfasigma, and speaker expenses from Alfasigma and Ferring Pharmaceuticals. N.P. has received research grant(s) from Bristol Myers Squibb, and reports personal fees from Takeda, Janssen, Pfizer, Bristol-Myers Squibb, Abbvie, Roche, Lilly, Allergan, and Celgene, outside the submitted work. N.P. has served as a speaker/advisory board member for Abbvie, Allergan, Bristol Myers Squibb, Celgene, Falk, Ferring, Janssen, Pfizer, Tillotts, Takeda and Vifor Pharma. Authors declare that they have no competing interests.

## Data and materials availability

The code used to generate the cytokine–cytokine interactions, figures, data and the subsequent analysis can be accessed in the project github repository (https://github.com/korcsmarosgroup/ulcerative-colitis-cytokine-networks). The generated cytokine networks can also be interactively accessed on the NDEx platform (96). RNA sequencing data has been deposited to [link].

## Notes

https://github.com/korcsmarosgroup/ulcerative-colitis-cytokine-networks

## References

1. M. J. Cameron, D. J. Kelvin, Cytokines, Chemokines and Their Receptors - Madame Curie Bioscience Database - NCBI Bookshelf (2013).

2. R. Morris, N. J. Kershaw, J. J. Babon, The molecular details of cytokine signaling via the JAK/STAT pathway. Protein Sci. 27, 1984–2009 (2018).

3. S. H. Chan, B. Perussia, J. W. Gupta, M. Kobayashi, M. Pospísil, H. A. Young, S. F. Wolf, D. Young, S. C. Clark, G. Trinchieri, Induction of interferon gamma production by natural killer cell stimulatory factor: characterization of the responder cells and synergy with other inducers. J. Exp. Med. 173, 869–879 (1991).

4. J. Ye, J. R. Ortaldo, K. Conlon, R. Winkler-Pickett, H. A. Young, Cellular and molecular mechanisms of IFN-gamma production induced by IL-2 and IL-12 in a human NK cell line. J. Leukoc. Biol. 58, 225–233 (1995).

5. J. H. Dufour, M. Dziejman, M. T. Liu, J. H. Leung, T. E. Lane, A. D. Luster, IFN-gamma-inducible protein 10 (IP-10; CXCL10)-deficient mice reveal a role for IP-10 in effector T cell generation and trafficking. J. Immunol. 168, 3195–3204 (2002).

6. M. Friedrich, M. Pohin, F. Powrie, Cytokine networks in the pathophysiology of inflammatory bowel disease. Immunity. 50, 992–1006 (2019).

7. S. C. Ng, H. Y. Shi, N. Hamidi, F. E. Underwood, W. Tang, E. I. Benchimol, R. Panaccione, S. Ghosh, J. C. Y. Wu, F. K. L. Chan, J. J. Y. Sung, G. G. Kaplan, Worldwide incidence and prevalence of inflammatory bowel disease in the 21st century: a systematic review of population-based studies. Lancet. 390, 2769–2778 (2017).

8. T. Kobayashi, B. Siegmund, C. Le Berre, S. C. Wei, M. Ferrante, B. Shen, C. N. Bernstein, S. Danese, L. Peyrin-Biroulet, T. Hibi, Ulcerative colitis. Nat. Rev. Dis. Primers. 6, 74 (2020).

9. T. Takeuchi, Cytokines and cytokine receptors as targets of immune-mediated inflammatory diseases-RA as a role model. Inflamm. Regen. 42, 35 (2022).

10. G. Schett, I. B. McInnes, M. F. Neurath, Reframing Immune-Mediated Inflammatory Diseases through Signature Cytokine Hubs. N. Engl. J. Med. 385, 628–639 (2021).

11. M. F. Neurath, Strategies for targeting cytokines in inflammatory bowel disease. Nat. Rev. Immunol. 24, 559–576 (2024).

12. M. F. Neurath, Current and emerging therapeutic targets for IBD. Nat. Rev. Gastroenterol. Hepatol. 14, 269–278 (2017).

13. D. Alsoud, B. Verstockt, C. Fiocchi, S. Vermeire, Breaking the therapeutic ceiling in drug development in ulcerative colitis. Lancet Gastroenterol. Hepatol. 6, 589–595 (2021).

14. N. R. West, A. N. Hegazy, B. M. J. Owens, S. J. Bullers, B. Linggi, S. Buonocore, M. Coccia, D. Görtz, S. This, K. Stockenhuber, J. Pott, M. Friedrich, G. Ryzhakov, F. Baribaud, C. Brodmerkel, C. Cieluch, N. Rahman, G. Müller-Newen, R. J. Owens, A. A. Kühl, F. Powrie, Oncostatin M drives intestinal inflammation and predicts response to tumor necrosis factor-neutralizing therapy in patients with inflammatory bowel disease. Nat. Med. 23, 579–589 (2017).

15. C. A. Lamb, A. Saifuddin, N. Powell, F. Rieder, The future of precision medicine to predict outcomes and control tissue remodeling in inflammatory bowel disease. Gastroenterology. 162, 1525–1542 (2022).

16. B. G. Feagan, B. E. Sands, W. J. Sandborn, M. Germinaro, M. Vetter, J. Shao, S. Sheng, J. Johanns, J. Panés, VEGA Study Group, Guselkumab plus golimumab combination therapy versus guselkumab or golimumab monotherapy in patients with ulcerative colitis (VEGA): a randomised, double-blind, controlled, phase 2, proof-of-concept trial. Lancet Gastroenterol. Hepatol. 8, 307–320 (2023).

17. G. Roda, M. Marocchi, A. Sartini, E. Roda, Cytokine networks in ulcerative colitis. Ulcers. 2011, 1–5 (2011).

18. M. Vebr, R. Pomahačová, J. Sýkora, J. Schwarz, A narrative review of cytokine networks: pathophysiological and therapeutic implications for inflammatory bowel disease pathogenesis. Biomedicines. 11 (2023), doi:10.3390/biomedicines11123229.

19. M. Olbei, J. P. Thomas, I. Hautefort, A. Treveil, B. Bohar, M. Madgwick, L. Gul, L. Csabai, D. Modos, T. Korcsmaros, CytokineLink: A Cytokine Communication Map to Analyse Immune Responses-Case Studies in Inflammatory Bowel Disease and COVID-19. Cells. 10 (2021), doi:10.3390/cells10092242.

20. J. E. Jansen, D. Aschenbrenner, H. H. Uhlig, M. C. Coles, E. A. Gaffney, A method for the inference of cytokine interaction networks. PLoS Comput. Biol. 18, e1010112 (2022).

21. H. Nie, P. Lin, Y. Zhang, Y. Wan, J. Li, C. Yin, L. Zhang, Single-cell meta-analysis of inflammatory bowel disease with scIBD. Nat. Comput. Sci. 3, 522–531 (2023).

22. R. Browaeys, W. Saelens, Y. Saeys, NicheNet: modeling intercellular communication by linking ligands to target genes. Nat. Methods. 17, 159–162 (2020).

23. A. Cui, T. Huang, S. Li, A. Ma, J. L. Pérez, C. Sander, D. B. Keskin, C. J. Wu, E. Fraenkel, N. Hacohen, Dictionary of immune responses to cytokines at single-cell resolution. Nature. 625, 377–384 (2024).

24. D. Masopust, C. P. Sivula, S. C. Jameson, Of mice, dirty mice, and men: using mice to understand human immunology. J. Immunol. 199, 383–388 (2017).

25. L. E. Wagar, R. M. DiFazio, M. M. Davis, Advanced model systems and tools for basic and translational human immunology. Genome Med. 10, 73 (2018).

26. J. Mestas, C. C. W. Hughes, Of mice and not men: differences between mouse and human immunology. J. Immunol. 172, 2731–2738 (2004).

27. K. Arnauts, B. Verstockt, A. S. Ramalho, S. Vermeire, C. Verfaillie, M. Ferrante, Ex vivo mimicking of inflammation in organoids derived from patients with ulcerative colitis. Gastroenterology. 159, 1564–1567 (2020).

28. V. Langer, E. Vivi, D. Regensburger, T. H. Winkler, M. J. Waldner, T. Rath, B. Schmid, L. Skottke, S. Lee, N. L. Jeon, T. Wohlfahrt, V. Kramer, P. Tripal, M. Schumann, S. Kersting, C. Handtrack, C. I. Geppert, K. Suchowski, R. H. Adams, C. Becker, M. Stürzl, IFN-γ drives inflammatory bowel disease pathogenesis through VE-cadherin-directed vascular barrier disruption. J. Clin. Invest. 129, 4691–4707 (2019).

29. N. C. Di Paolo, D. M. Shayakhmetov, Interleukin 1α and the inflammatory process. Nat. Immunol. 17, 906–913 (2016).

30. C. P. McEntee, C. M. Finlay, E. C. Lavelle, Divergent Roles for the IL-1 Family in Gastrointestinal Homeostasis and Inflammation. Front. Immunol. 10, 1266 (2019).

31. A. Konrad, M. Lehrke, V. Schachinger, F. Seibold, R. Stark, T. Ochsenkühn, K. G. Parhofer, B. Göke, U. C. Broedl, Resistin is an inflammatory marker of inflammatory bowel disease in humans. Eur. J. Gastroenterol. Hepatol. 19, 1070–1074 (2007).

32. N. Silswal, A. K. Singh, B. Aruna, S. Mukhopadhyay, S. Ghosh, N. Z. Ehtesham, Human resistin stimulates the pro-inflammatory cytokines TNF-alpha and IL-12 in macrophages by NF-kappaB-dependent pathway. Biochem. Biophys. Res. Commun. 334, 1092–1101 (2005).

33. C. Guo, K. Wu, X. Liang, Y. Liang, R. Li, Infliximab clinically treating ulcerative colitis: A systematic review and meta-analysis. Pharmacol. Res. 148, 104455 (2019).

34. A. Andoh, A. Nishida, Molecular basis of intestinal fibrosis in inflammatory bowel disease. Inflamm. Intest. Dis. 7, 119–127 (2023).

35. C. Eftychi, R. Schwarzer, K. Vlantis, L. Wachsmuth, M. Basic, P. Wagle, M. F. Neurath, C. Becker, A. Bleich, M. Pasparakis, Temporally Distinct Functions of the Cytokines IL-12 and IL-23 Drive Chronic Colon Inflammation in Response to Intestinal Barrier Impairment. Immunity. 51, 367–380.e4 (2019).

36. I. H. Goenawan, K. Bryan, D. J. Lynn, DyNet: visualization and analysis of dynamic molecular interaction networks. Bioinformatics. 32, 2713–2715 (2016).

37. P. Pavlidis, A. Tsakmaki, E. Pantazi, K. Li, D. Cozzetto, J. Digby-Bell, F. Yang, J. W. Lo, E. Alberts, A. C. C. Sa, U. Niazi, J. Friedman, A. K. Long, Y. Ding, C. D. Carey, C. Lamb, M. Saqi, M. Madgwick, L. Gul, A. Treveil, N. Powell, Interleukin-22 regulates neutrophil recruitment in ulcerative colitis and is associated with resistance to ustekinumab therapy. Nat. Commun. 13, 5820 (2022).

38. C. A. Lindemans, M. Calafiore, A. M. Mertelsmann, M. H. O’Connor, J. A. Dudakov, R. R. Jenq, E. Velardi, L. F. Young, O. M. Smith, G. Lawrence, J. A. Ivanov, Y.-Y. Fu, S. Takashima, G. Hua, M. L. Martin, K. P. O’Rourke, Y.-H. Lo, M. Mokry, M. Romera-Hernandez, T. Cupedo, A. M. Hanash, Interleukin-22 promotes intestinal-stem-cell-mediated epithelial regeneration. Nature. 528, 560–564 (2015).

39. D. Bauché, B. Joyce-Shaikh, J. Fong, A. V. Villarino, K. S. Ku, R. Jain, Y.-C. Lee, L. Annamalai, J. H. Yearley, D. J. Cua, IL-23 and IL-2 activation of STAT5 is required for optimal IL-22 production in ILC3s during colitis. Sci. Immunol. 5 (2020), doi:10.1126/sciimmunol.aav1080.

40. S. Verstockt, B. Verstockt, K. Machiels, M. Vancamelbeke, M. Ferrante, I. Cleynen, G. De Hertogh, S. Vermeire, Oncostatin M is a biomarker of diagnosis, worse disease prognosis, and therapeutic nonresponse in inflammatory bowel disease. Inflamm. Bowel Dis. 27, 1564–1575 (2021).

41. X. Zhao, W. Yang, T. Yu, Y. Yu, X. Cui, Z. Zhou, H. Yang, Y. Yu, A. J. Bilotta, S. Yao, J. Xu, J. Zhou, G. S. Yochum, W. A. Koltun, A. Portolese, D. Zeng, J. Xie, I. V. Pinchuk, H. Zhang, Y. Cong, Th17 Cell-Derived Amphiregulin Promotes Colitis-Associated Intestinal Fibrosis Through Activation of mTOR and MEK in Intestinal Myofibroblasts. Gastroenterology. 164, 89–102 (2023).

42. M. Kurimoto, T. Watanabe, K. Kamata, K. Minaga, M. Kudo, IL-33 as a Critical Cytokine for Inflammation and Fibrosis in Inflammatory Bowel Diseases and Pancreatitis. Front. Physiol. 12, 781012 (2021).

43. M. A. Williams, A. O’Callaghan, S. C. Corr, IL-33 and IL-18 in Inflammatory Bowel Disease Etiology and Microbial Interactions. Front. Immunol. 10, 1091 (2019).

44. B. Siegmund, G. Fantuzzi, F. Rieder, F. Gamboni-Robertson, H. A. Lehr, G. Hartmann, C. A. Dinarello, S. Endres, A. Eigler, Neutralization of interleukin-18 reduces severity in murine colitis and intestinal IFN-gamma and TNF-alpha production. Am. J. Physiol. Regul. Integr. Comp. Physiol. 281, R1264–73 (2001).

45. D. Polosukhina, K. Singh, M. Asim, D. P. Barry, M. M. Allaman, D. M. Hardbower, M. B. Piazuelo, M. K. Washington, A. P. Gobert, K. T. Wilson, L. A. Coburn, CCL11 exacerbates colitis and inflammation-associated colon tumorigenesis. Oncogene. 40, 6540–6546 (2021).

46. T. Castro-Dopico, A. Fleming, T. W. Dennison, J. R. Ferdinand, K. Harcourt, B. J. Stewart, Z. Cader, Z. K. Tuong, C. Jing, L. S. C. Lok, R. J. Mathews, A. Portet, A. Kaser, S. Clare, M. R. Clatworthy, GM-CSF Calibrates Macrophage Defense and Wound Healing Programs during Intestinal Infection and Inflammation. Cell Rep. 32, 107857 (2020).

47. E. Landy, H. Carol, A. Ring, S. Canna, Biological and clinical roles of IL-18 in inflammatory diseases. Nat. Rev. Rheumatol. 20, 33–47 (2024).

48. L. Chen, X.-L. Zhong, W.-Y. Cao, M.-L. Mao, D.-D. Liu, W.-J. Liu, X.-Y. Zu, J.-H. Liu, IGF2/IGF2R/Sting signaling as a therapeutic target in DSS-induced ulcerative colitis. Eur. J. Pharmacol. 960, 176122 (2023).

49. Z. Xie, G. Zhou, M. Zhang, J. Han, Y. Wang, X. Li, Q. Wu, M. Li, S. Zhang, Recent developments on BMPs and their antagonists in inflammatory bowel diseases. Cell Death Discov. 9, 210 (2023).

50. M. Sochal, M. Ditmer, A. Gabryelska, P. Białasiewicz, The Role of Brain-Derived Neurotrophic Factor in Immune-Related Diseases: A Narrative Review. J. Clin. Med. 11 (2022), doi:10.3390/jcm11206023.

51. R. Hjortebjerg, K. L. Thomsen, J. Agnholt, J. Frystyk, The IGF system in patients with inflammatory bowel disease treated with prednisolone or infliximab: potential role of the stanniocalcin-2 / PAPP-A / IGFBP-4 axis. BMC Gastroenterol. 19, 83 (2019).

52. X. Hu, J. Li, M. Fu, X. Zhao, W. Wang, The JAK/STAT signaling pathway: from bench to clinic. Signal Transduct. Target. Ther. 6, 402 (2021).

53. S. Honap, A. Agorogianni, M. J. Colwill, S. K. Mehta, F. Donovan, R. Pollok, A. Poullis, K. Patel, JAK inhibitors for inflammatory bowel disease: recent advances. Frontline Gastroenterol. 15, 59–69 (2024).

54. W. J. Sandborn, C. Su, B. E. Sands, G. R. D’Haens, S. Vermeire, S. Schreiber, S. Danese, B. G. Feagan, W. Reinisch, W. Niezychowski, G. Friedman, N. Lawendy, D. Yu, D. Woodworth, A. Mukherjee, H. Zhang, P. Healey, J. Panés, OCTAVE Induction 1, OCTAVE Induction 2, and OCTAVE Sustain Investigators, Tofacitinib as induction and maintenance therapy for ulcerative colitis. N. Engl. J. Med. 376, 1723–1736 (2017).

55. S. Danese, S. Vermeire, W. Zhou, A. L. Pangan, J. Siffledeen, S. Greenbloom, X. Hébuterne, G. D’Haens, H. Nakase, J. Panés, P. D. R. Higgins, P. Juillerat, J. O. Lindsay, E. V. Loftus, W. J. Sandborn, W. Reinisch, M.-H. Chen, Y. Sanchez Gonzalez, B. Huang, W. Xie, R. Panaccione, Upadacitinib as induction and maintenance therapy for moderately to severely active ulcerative colitis: results from three phase 3, multicentre, double-blind, randomised trials. Lancet. 399, 2113–2128 (2022).

56. D. Ochoa, A. Hercules, M. Carmona, D. Suveges, J. Baker, C. Malangone, I. Lopez, A. Miranda, C. Cruz-Castillo, L. Fumis, M. Bernal-Llinares, K. Tsukanov, H. Cornu, K. Tsirigos, O. Razuvayevskaya, A. Buniello, J. Schwartzentruber, M. Karim, B. Ariano, R. E. Martinez Osorio, E. M. McDonagh, The next-generation Open Targets Platform: reimagined, redesigned, rebuilt. Nucleic Acids Res. 51, D1353–D1359 (2023).

57. R. Thiébaut, S. Kotti, C. Jung, F. Merlin, J. F. Colombel, M. Lemann, S. Almer, C. Tysk, M. O’Morain, M. Gassull, V. Binder, Y. Finkel, L. Pascoe, J. P. Hugot, TNFSF15 polymorphisms are associated with susceptibility to inflammatory bowel disease in a new European cohort. Am. J. Gastroenterol. 104, 384–391 (2009).

58. F. Furfaro, L. Alfarone, D. Gilardi, C. Correale, M. Allocca, G. Fiorino, M. Argollo, A. Zilli, E. Zacharopoulou, L. Loy, G. Roda, S. Danese, TL1A: A new potential target in the treatment of inflammatory bowel disease. Curr. Drug Targets. 22, 760–769 (2021).

59. V. Solitano, V. Jairath, F. Ungaro, L. Peyrin-Biroulet, S. Danese, TL1A inhibition for inflammatory bowel disease treatment: From inflammation to fibrosis. MED. 5, 386–400 (2024).

60. S. Jin, J. Chin, S. Seeber, J. Niewoehner, B. Weiser, N. Beaucamp, J. Woods, C. Murphy, A. Fanning, F. Shanahan, K. Nally, R. Kajekar, A. Salas, N. Planell, J. Lozano, J. Panes, H. Parmar, J. DeMartino, S. Narula, D. A. Thomas-Karyat, TL1A/TNFSF15 directly induces proinflammatory cytokines, including TNFα, from CD3+CD161+ T cells to exacerbate gut inflammation. Mucosal Immunol. 6, 886–899 (2013).

61. V. Valatas, G. Kolios, G. Bamias, TL1A (TNFSF15) and DR3 (TNFRSF25): A Co-stimulatory System of Cytokines With Diverse Functions in Gut Mucosal Immunity. Front. Immunol. 10, 583 (2019).

62. M. Hassan-Zahraee, Z. Ye, L. Xi, M. L. Baniecki, X. Li, C. L. Hyde, J. Zhang, N. Raha, F. Karlsson, J. Quan, D. Ziemek, S. Neelakantan, C. Lepsy, J. R. Allegretti, J. Romatowski, E. J. Scherl, M. Klopocka, S. Danese, D. E. Chandra, U. Schoenbeck, K. E. Hung, Antitumor Necrosis Factor-like Ligand 1A Therapy Targets Tissue Inflammation and Fibrosis Pathways and Reduces Gut Pathobionts in Ulcerative Colitis. Inflamm. Bowel Dis. 28, 434–446 (2022).

63. S. Kany, J. T. Vollrath, B. Relja, Cytokines in inflammatory disease. Int. J. Mol. Sci. 20 (2019), doi:10.3390/ijms20236008.

64. I. F. Charo, R. M. Ransohoff, The many roles of chemokines and chemokine receptors in inflammation. N. Engl. J. Med. 354, 610–621 (2006).

65. P. A. Morel, R. E. C. Lee, J. R. Faeder, Demystifying the cytokine network: Mathematical models point the way. Cytokine. 98, 115–123 (2017).

66. I. B. McInnes, E. M. Gravallese, Immune-mediated inflammatory disease therapeutics: past, present and future. Nat. Rev. Immunol. 21, 680–686 (2021).

67. K. Kveler, E. Starosvetsky, A. Ziv-Kenet, Y. Kalugny, Y. Gorelik, G. Shalev-Malul, N. Aizenbud-Reshef, T. Dubovik, M. Briller, J. Campbell, J. C. Rieckmann, N. Asbeh, D. Rimar, F. Meissner, J. Wiser, S. S. Shen-Orr, Immune-centric network of cytokines and cells in disease context identified by computational mining of PubMed. Nat. Biotechnol. 36, 651–659 (2018).

68. M. B. Atallah, V. Tandon, K. J. Hiam, H. Boyce, M. Hori, W. Atallah, M. H. Spitzer, E. Engleman, P. Mallick, ImmunoGlobe: enabling systems immunology with a manually curated intercellular immune interaction network. BMC Bioinformatics. 21, 346 (2020).

69. K. Sommer, M. Wiendl, T. M. Müller, K. Heidbreder, C. Voskens, M. F. Neurath, S. Zundler, Intestinal Mucosal Wound Healing and Barrier Integrity in IBD-Crosstalk and Trafficking of Cellular Players. Front Med (Lausanne). 8, 643973 (2021).

70. E. Bartee, G. McFadden, Cytokine synergy: an underappreciated contributor to innate anti-viral immunity. Cytokine. 63, 237–240 (2013).

71. E. Armingol, A. Officer, O. Harismendy, N. E. Lewis, Deciphering cell–cell interactions and communication from gene expression. Nat. Rev. Genet. (2020), doi:10.1038/s41576-020-00292-x.

72. P. S. L. Schäfer, D. Dimitrov, E. J. Villablanca, J. Saez-Rodriguez, Integrating single-cell multi-omics and prior biological knowledge for a functional characterization of the immune system. Nat. Immunol. 25, 405–417 (2024).

73. H. Daneshpour, H. Youk, Modeling cell-cell communication for immune systems across space and time. Current Opinion in Systems Biology. 18, 44–52 (2019).

74. C. G. Williams, H. J. Lee, T. Asatsuma, R. Vento-Tormo, A. Haque, An introduction to spatial transcriptomics for biomedical research. Genome Med. 14, 68 (2022).

75. I. Kleino, P. Frolovaitė, T. Suomi, L. L. Elo, Computational solutions for spatial transcriptomics. Comput. Struct. Biotechnol. J. 20, 4870–4884 (2022).

76. E. Armingol, A. Ghaddar, C. J. Joshi, H. Baghdassarian, I. Shamie, J. Chan, H.-L. Her, S. Berhanu, A. Dar, F. Rodriguez-Armstrong, O. Yang, E. J. O’Rourke, N. E. Lewis, Inferring a spatial code of cell-cell interactions across a whole animal body. PLoS Comput. Biol. 18, e1010715 (2022).

77. G. Palla, D. S. Fischer, A. Regev, F. J. Theis, Spatial components of molecular tissue biology. Nat. Biotechnol. 40, 308–318 (2022).

78. I. Hautefort, M. Poletti, D. Papp, T. Korcsmaros, Everything You Always Wanted to Know About Organoid-Based Models (and Never Dared to Ask). Cell. Mol. Gastroenterol. Hepatol. 14, 311–331 (2022).

79. D. Papp, T. Korcsmaros, I. Hautefort, Revolutionising immune research with organoid-based co-culture and chip systems. Clin. Exp. Immunol. (2024), doi:10.1093/cei/uxae004.

80. T. Recaldin, L. Steinacher, B. Gjeta, M. F. Harter, L. Adam, K. Kromer, M. P. Mendes, M. Bellavista, M. Nikolaev, G. Lazzaroni, R. Krese, U. Kilik, D. Popovic, B. Stoll, R. Gerard, M. Bscheider, M. Bickle, L. Cabon, J. G. Camp, N. Gjorevski, Human organoids with an autologous tissue-resident immune compartment. Nature (2024), doi:10.1038/s41586-024-07791-5.

81. J.-P. Villemin, L. Bassaganyas, D. Pourquier, F. Boissière, S. Cabello-Aguilar, E. Crapez, R. Tanos, E. Cornillot, A. Turtoi, J. Colinge, Inferring ligand-receptor cellular networks from bulk and spatial transcriptomic datasets with BulkSignalR. Nucleic Acids Res. 51, 4726–4744 (2023).

82. Y. Hao, T. Stuart, M. H. Kowalski, S. Choudhary, P. Hoffman, A. Hartman, A. Srivastava, G. Molla, S. Madad, C. Fernandez-Granda, R. Satija, Dictionary learning for integrative, multimodal and scalable single-cell analysis. Nat. Biotechnol. 42, 293–304 (2024).

83. S. Mangiola, M. A. Doyle, A. T. Papenfuss, Interfacing Seurat with the R tidy universe. Bioinformatics. 37, 4100–4107 (2021).

84. J. Kinchen, H. H. Chen, K. Parikh, A. Antanaviciute, M. Jagielowicz, D. Fawkner-Corbett, N. Ashley, L. Cubitt, E. Mellado-Gomez, M. Attar, E. Sharma, Q. Wills, R. Bowden, F. C. Richter, D. Ahern, K. D. Puri, J. Henault, F. Gervais, H. Koohy, A. Simmons, Structural remodeling of the human colonic mesenchyme in inflammatory bowel disease. Cell. 175, 372–386.e17 (2018).

85. K. Parikh, A. Antanaviciute, D. Fawkner-Corbett, M. Jagielowicz, A. Aulicino, C. Lagerholm, S. Davis, J. Kinchen, H. H. Chen, N. K. Alham, N. Ashley, E. Johnson, P. Hublitz, L. Bao, J. Lukomska, R. S. Andev, E. Björklund, B. M. Kessler, R. Fischer, R. Goldin, A. Simmons, Colonic epithelial cell diversity in health and inflammatory bowel disease. Nature. 567, 49–55 (2019).

86. D. Corridoni, A. Antanaviciute, T. Gupta, D. Fawkner-Corbett, A. Aulicino, M. Jagielowicz, K. Parikh, E. Repapi, S. Taylor, D. Ishikawa, R. Hatano, T. Yamada, W. Xin, H. Slawinski, R. Bowden, G. Napolitani, O. Brain, C. Morimoto, H. Koohy, A. Simmons, Single-cell atlas of colonic CD8+ T cells in ulcerative colitis. Nat. Med. 26, 1480–1490 (2020).

87. P. Shannon, A. Markiel, O. Ozier, N. S. Baliga, J. T. Wang, D. Ramage, N. Amin, B. Schwikowski, T. Ideker, Cytoscape: a software environment for integrated models of biomolecular interaction networks. Genome Res. 13, 2498–2504 (2003).

88. E. Paradis, J. Claude, K. Strimmer, APE: Analyses of Phylogenetics and Evolution in R language. Bioinformatics. 20, 289–290 (2004).

89. I. Letunic, P. Bork, Interactive Tree Of Life (iTOL) v5: an online tool for phylogenetic tree display and annotation. Nucleic Acids Res. 49, W293–W296 (2021).

90. D. P. Gil, J. N. Law, T. M. Murali, The PathLinker app: Connect the dots in protein interaction networks. [version 1; peer review: 1 approved, 2 approved with reservations]. F1000Res. 6, 58 (2017).

91. Andrews S., Babraham Bioinformatics - FastQC A Quality Control tool for High Throughput Sequence Data. Babraham Bioinformatics (2010), (available at https://www.bioinformatics.babraham.ac.uk/projects/fastqc/).

92. A. Dobin, C. A. Davis, F. Schlesinger, J. Drenkow, C. Zaleski, S. Jha, P. Batut, M. Chaisson, T. R. Gingeras, STAR: ultrafast universal RNA-seq aligner. Bioinformatics. 29, 15–21 (2013).

93. Y. Liao, G. K. Smyth, W. Shi, featureCounts: an efficient general purpose program for assigning sequence reads to genomic features. Bioinformatics. 30, 923–930 (2014).

94. P. Ewels, M. Magnusson, S. Lundin, M. Käller, MultiQC: summarize analysis results for multiple tools and samples in a single report. Bioinformatics. 32, 3047–3048 (2016).

95. M. I. Love, W. Huber, S. Anders, Moderated estimation of fold change and dispersion for RNA-seq data with DESeq2. Genome Biol. 15, 550 (2014).

96. D. Pratt, J. Chen, R. Pillich, V. Rynkov, A. Gary, B. Demchak, T. Ideker, Ndex 2.0: A clearinghouse for research on cancer pathways. Cancer Res. 77, e58–e61 (2017).

