## Supplementary materials for "Decoding Cytokine Networks in Ulcerative Colitis to Identify Pathogenic Mechanisms and Therapeutic Targets"

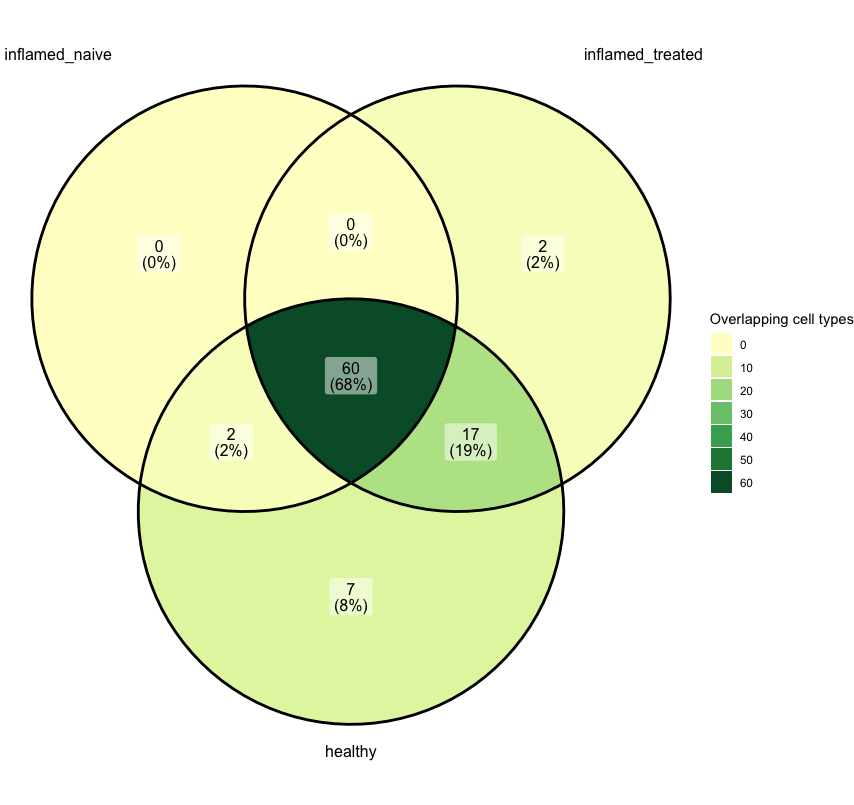

Figure S1. Number of overlapping cell types between the inflamed conditions and healthy.

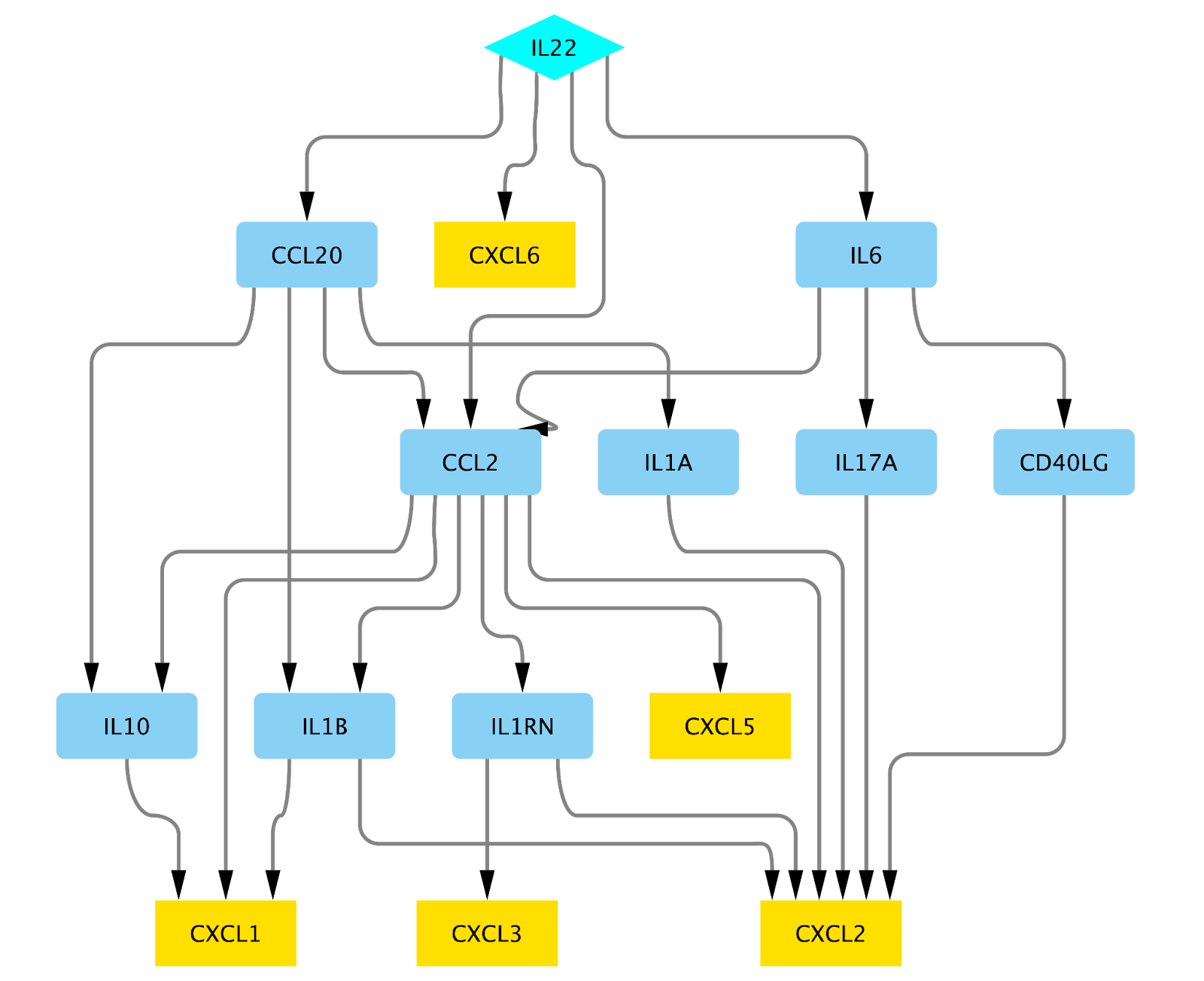

Figure S2. IL22 signalling to CXC-family cytokines.

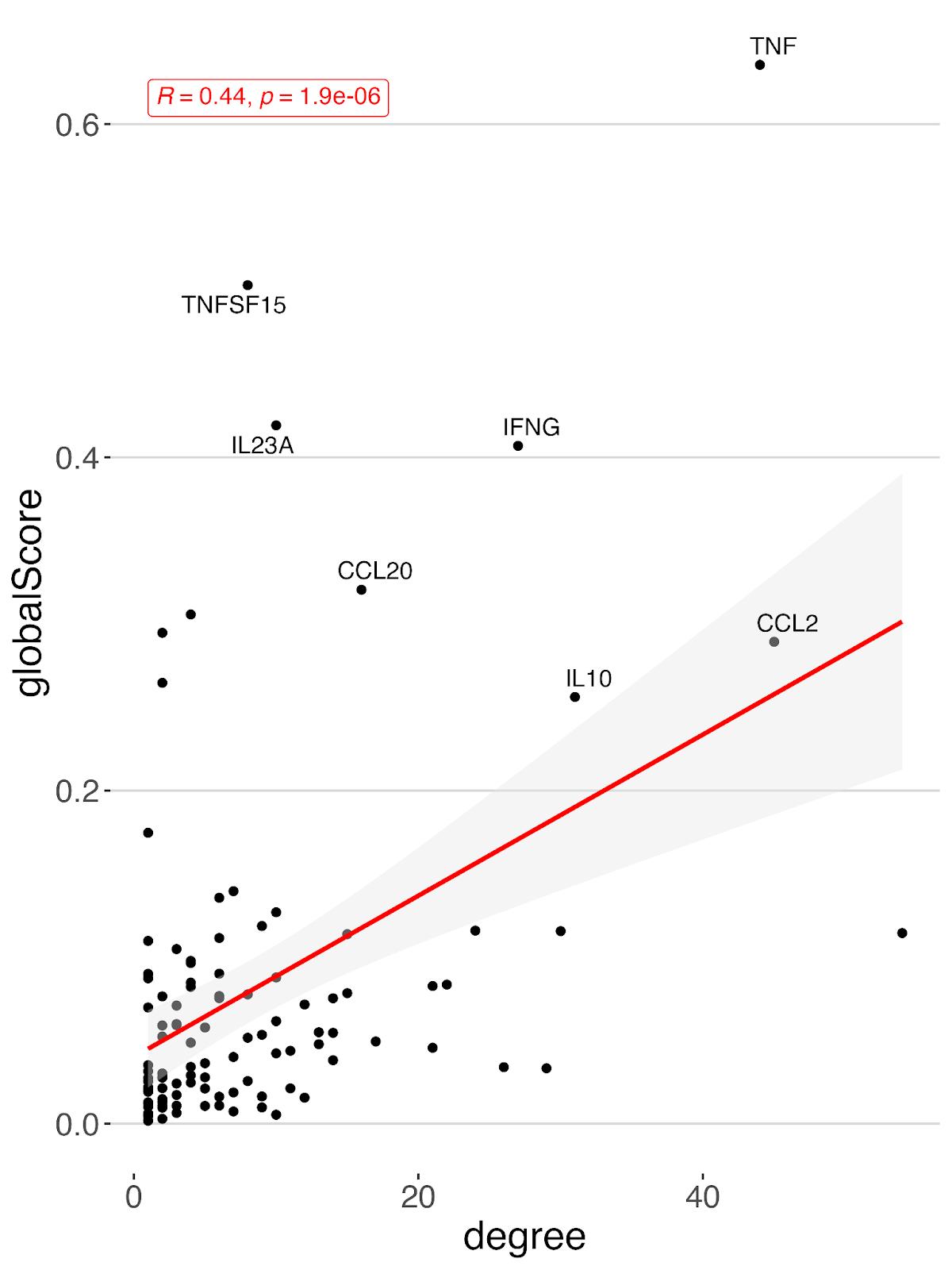

Figure S3. Correlation of OpenTargets global score with degree centrality.

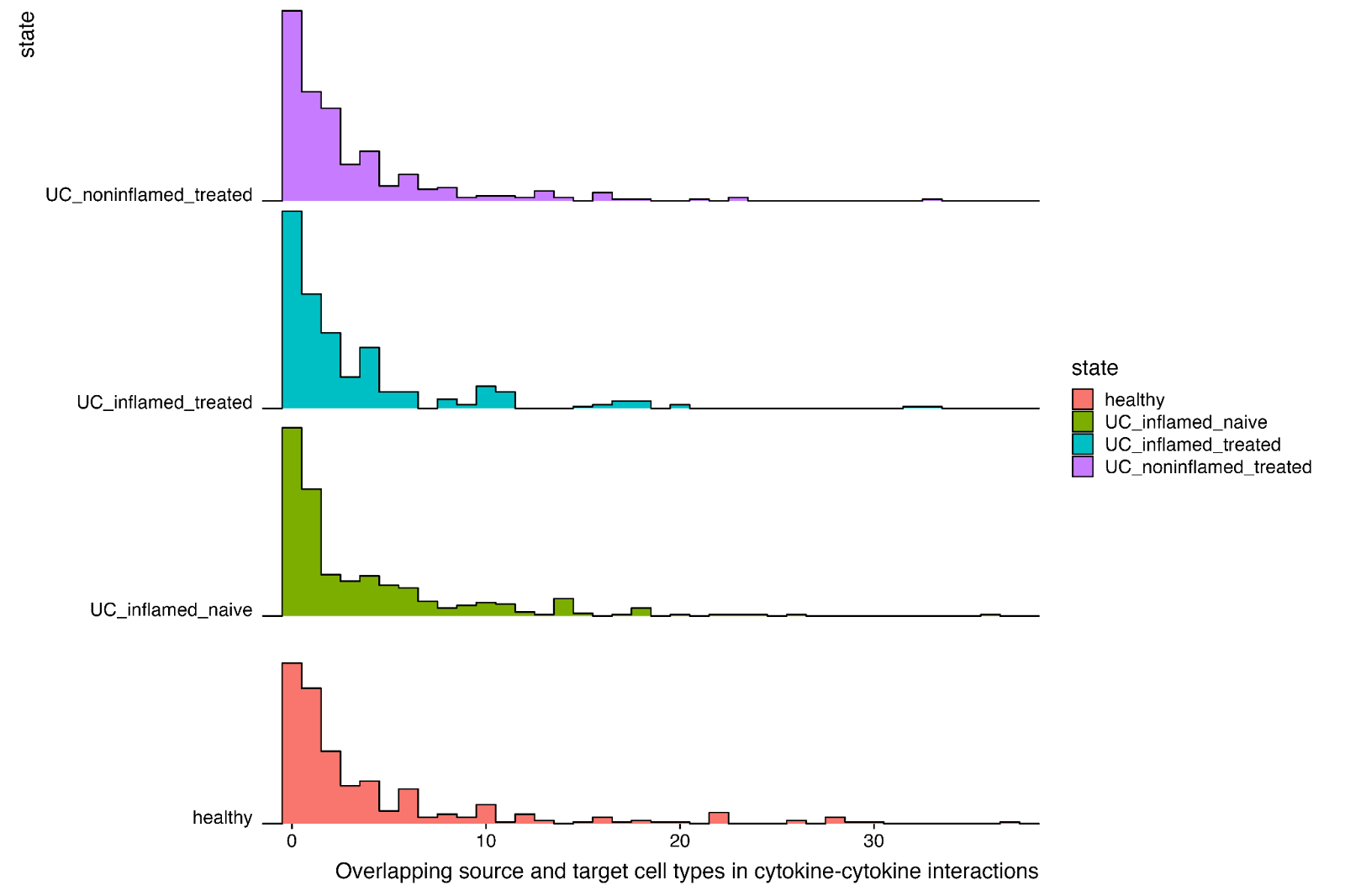

Figure S4. Distribution of the number of overlapping source and target cell types in cytokine–cytokine interactions.

Supplementary Table 1: List of cytokines in the cytokine networks

Supplementary Table 1: List of cytokines in the cytokine networks

**Cytokines**

ADM

ANGPTL4

BMP6

BMP7

CCL1

CCL11

CCL13

CCL14

CCL15

CCL17

CCL18

CCL2

CCL20

CCL21

CCL26

CCL3

CCL4

CCL5

CCL7

CCL8

CD40LG

CSF2

CX3CL1

CXCL10

CXCL11

CXCL13

CXCL14

CXCL9

FGF2

FGF7

HGF

IFNG

IGF2

IL10

IL15

IL16

IL17A

IL17B

IL18

IL1A

IL1B

IL1RN

IL22

IL23A

IL33

IL6

IL7

KITLG

MIF

OSM

PDGFB

PDGFC

PDGFD

PF4

RETN

SPP1

TGFB1

TNF

TNFSF11

TNFSF12

TNFSF14

TNFSF15

TSLP

WNT5A

HBEGF

TGFB2

VEGFA

CCL16

FGF23

BMP2

LIF

BDNF

CXCL2

IL2

IL4

BMP4

GDF15

AREG

CXCL1

CXCL5

EBI3

CD70

IGF1

IL12A

TNFSF13

TNFSF13B

TNFSF8

CXCL3

ANGPT1

MST1

PDGFA

CSF1

CCL23

IL19

IL32

LTA

CXCL12

CXCL6

IL11

NGF

IL13

CSF3

LTB

IL17F

IL24

CCL19

CCL24

IL37

TNFSF4

VEGFC

CCL22

CXCL16
